# Cross-tissue immune cell analysis reveals tissue-specific adaptations and clonal architecture in humans

**DOI:** 10.1101/2021.04.28.441762

**Authors:** C Domínguez Conde, C Xu, LB Jarvis, T Gomes, SK Howlett, DB Rainbow, O Suchanek, HW King, L Mamanova, K Polanski, N Huang, ES Fasouli, KT Mahbubani, M Prete, L Tuck, N Richoz, ZK Tuong, L Campos, HS Mousa, EJ Needham, S Pritchard, T Li, R Elmentaite, J Park, DK Menon, OA Bayraktar, LK James, KB Meyer, MR Clatworthy, K Saeb-Parsy, JL Jones, SA Teichmann

## Abstract

Despite their crucial role in health and disease, our knowledge of immune cells within human tissues remains limited. Here, we surveyed the immune compartment of 15 tissues of six deceased adult donors by single-cell RNA sequencing and paired VDJ sequencing. To systematically resolve immune cell heterogeneity across tissues, we developed CellTypist, a machine learning tool for rapid and precise cell type annotation. Using this approach, combined with detailed curation, we determined the tissue distribution of 45 finely phenotyped immune cell types and states, revealing hitherto unappreciated tissue-specific features and clonal architecture of T and B cells. In summary, our multi-tissue approach lays the foundation for identifying highly resolved immune cell types by leveraging a common reference dataset, tissue-integrated expression analysis and antigen receptor sequencing.

**One Sentence Summary:** We provide an immune cell atlas, including antigen receptor repertoire profiling, across lymphoid and non-lymphoid human tissues.

## Main Text

The immune system is a dynamic and integrated network made up of many different cell types distributed across the entire organism that act together to ensure effective host defence. In recent years, a growing appreciation of immune ontogeny and diversity across tissues has emerged. For example, we have gained new insights into how macrophages derived in embryogenesis contribute to the unique adaptation of tissue-resident myeloid cells, such as Langerhans cells in the skin, microglia in the brain and Kupffer cells in the liver (*1*–*3*). Other populations, such as innate lymphoid cells (ILCs), including natural killer (NK) cells and non-conventional (NKT, MAIT and γδ) T cells, have circulating counterparts but are highly enriched at barrier/mucosal sites where they mediate tissue defence and repair (*4*). In addition, following resolution of an immune response, antigen-specific, long-lived tissue-resident memory T cells (TRMs) have been shown to persist in many tissues, where they provide a first line of defence against secondary infections (reviewed in (*5*–*7*)).

Despite the central role of tissue immunity in homeostasis and disease, much of our current knowledge comes from animal studies. Historically, human immune cell analysis has focused on peripheral blood, providing a biased and incomplete view of human immunity. Recent work interrogating human tissue immunity using flow cytometry (*8*–*11*) and several organ-focused single-cell RNA sequencing (scRNA-seq) studies (*12*–*16*) have begun to address this knowledge gap, but few have analysed immune cells across multiple tissues from the same individual, which allows direct and controlled comparisons to be made. One such study by Szabo *et al*. has reported an analysis of T cells in three tissues from two donors (*17*). However, despite the effort to assemble murine (*18*) and human *(19, 20)* multi-tissue atlases, large-scale cross-tissue scRNA-seq studies that investigate tissue-specific features of the immune compartment remain scarce. To address this, here, we comprehensively profiled immune cell populations isolated from 15 donor-matched tissues from six deceased individuals representing in the order of 100,000 single cell transcriptomes.

Annotation of increasingly large single cell transcriptomics data sets remains a challenge, including the identification of rare cell subsets, and distinguishing previously described cell populations from novel discoveries. To capture the vast cellular diversity for annotating muti-tissue immune cells, we developed CellTypist, a machine learning framework for cell type prediction compiled from published studies across 20 human tissues. Combining this approach with the in-depth dissection of cellular heterogeneity within the myeloid and lymphoid compartments, we determined the frequency and transcriptional characteristics of immune cell types and states across human tissues. Moreover, we inferred the patterns of T and B cell migration across different tissues and their transition between cell states on the basis of antigen-receptor identity, gaining insights into tissue-dependent phenotypic and clonal plasticity.

## Results

### Single cell transcriptomics and antigen receptor sequence analysis of tissue mononuclear cells

To systematically assess immune cell type heterogeneity across human tissues, we performed scRNA-seq on 15 different tissues from six deceased organ donors (**Fig. 1A,B, Supplementary Table 1)**. Briefly, cells were isolated using a uniform protocol across tissues, with the exception of blood and bone marrow samples (see Methods for details). The tissues studied included primary (thymus and bone marrow) and secondary (spleen, thoracic and mesenteric lymph nodes) lymphoid organs, mucosal tissues (gut and lung), as well as liver, skeletal muscle and omentum. After stringent quality control, we obtained a total of 78,844 high quality hematopoietic cells, with higher yields from the lymphoid tissues in all donors (**Fig. S1A**,**B**). Using manual annotation based on marker gene expression (**Fig. S1C**), we identified 14 major cell populations represented across all donors and tissues from three major immune compartments: (i) T cells and ILCs, (ii) B cells and plasma cells, and (iii) myeloid cells (**Fig. 1C,D, Fig. S1D**,**E**). In addition, we identified four small distinct clusters consisting of mast cells, megakaryocytes/platelets (Mgk), plasmacytoid dendritic cells (pDCs) and a small population of immune progenitor cells. As expected, progenitors and megakaryocyte clusters were primarily found in the bone marrow and blood; macrophage and mast cell populations were enriched in the lung and lymphocytes mainly came from lymphoid organs (**Fig. S1E**). Analyses of the cell type composition within each tissue (**Fig. S1F**) further revealed that lymph nodes and thymus were rich in CD4+ T cells, while in the liver, spleen, bone marrow and gut, CD8+ T cells were predominant. In addition, gut regions were abundant in plasma cells, and the immune compartment of the lung parenchyma was dominated by monocytes and macrophages.

**Fig. 1.**
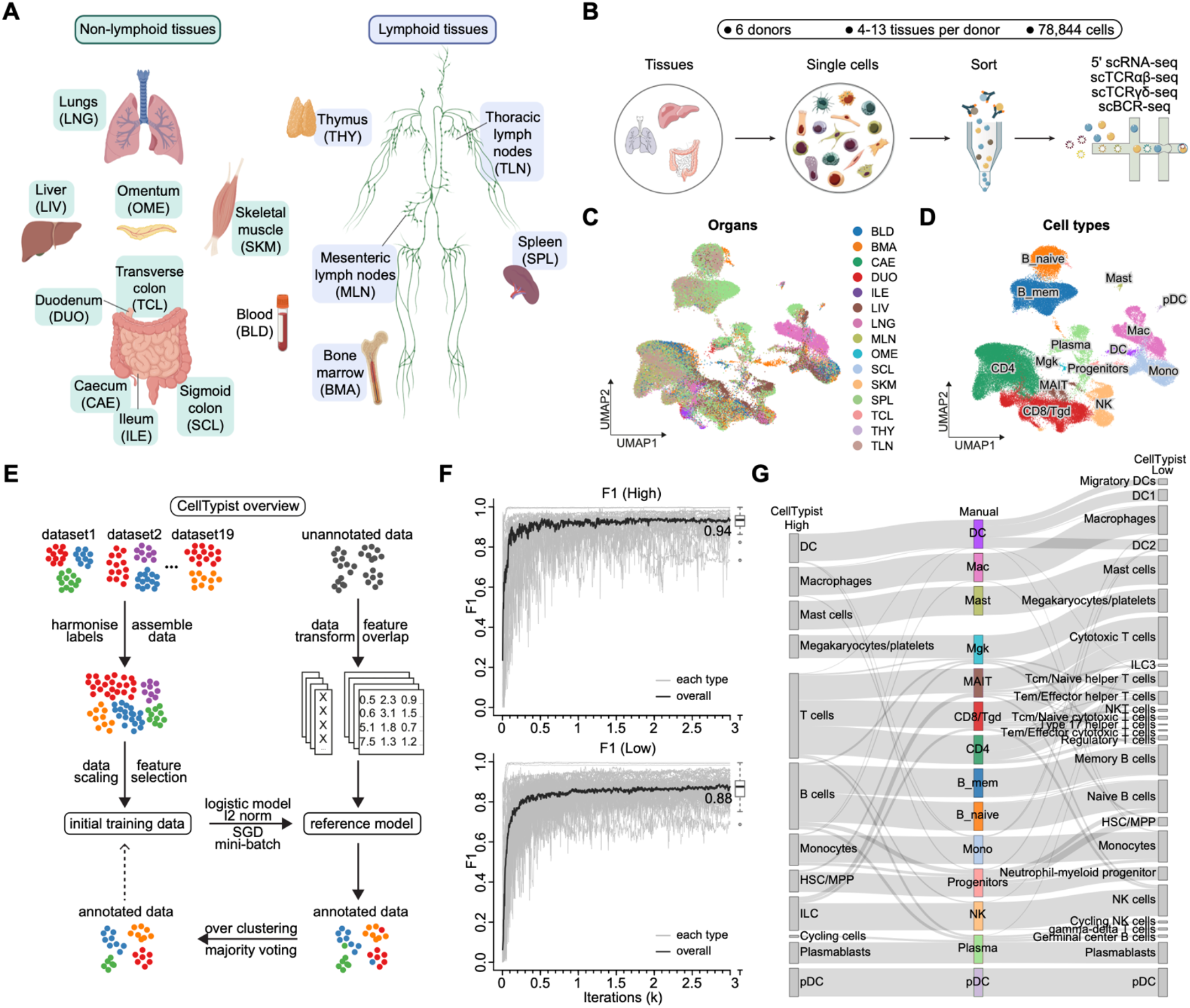
Automated annotation of immune cells across human tissues using CellTypist. (**A**) Schematic showing sample collections of human lymphoid and non-lymphoid tissues and their assigned tissue name acronyms. (**B**) Schematic of single-cell transcriptome profiling and paired sequencing of αβ TCR, γδ TCR and BCR variable regions. (**C**) UMAP visualization of the immune cell compartment colored by tissues. (**D**) As in (**C**), but colored by manually annotated cell types. (**E**) Workflow of CellTypist including data collection, processing, model training and cell type prediction. (**F**) Performance curves showing the F1 score at each iteration of training using mini-batch stochastic gradient descent for high- and low-hierarchy CellTypist models, respectively. The black curve represents the median F1 score averaged across the individual F1 scores of all predicted cell types (grey curves). (**G**) Sankey plot showing the fractions of CellTypist high-hierarchy model-derived labels (left) as compared to the manually defined clusters (center), as well as the fractions of CellTypist low-hierarchy model-derived labels (right).

For further analysis of the identity of cell states, we developed a machine learning framework trained on published datasets from 20 tissues, as described in the next section.

### CellTypist: a novel tool for annotating immune cell populations across tissues

Robust cell type annotation remains a major challenge in single-cell transcriptomics. To address the cellular heterogeneity in our cross-tissue study, we developed CellTypist (https://github.com/Teichlab/celltypist), a lightweight and interpretable cell type classification pipeline built on an expandable cell type reference atlas assembled from multiple tissues and studies (**Fig. 1E, Supplementary Note**). In brief, CellTypist is currently based on 20 tissues from 19 reference datasets (**Supplementary Note Figure 1A**) where cell type labels are harmonized at different hierarchically-structured levels (**Supplementary Note Figure 1B**). This hierarchy is of particular importance for determining immune cell identity at different levels of specificity (**Supplementary Note Table 1**), and is later implemented under a fine-tuned logistic regression framework using stochastic gradient descent learning. Performance of the derived models, as measured by precision, recall and global F1-score, reached ∼0.9 for cell type classifications at both the high- and low-hierarchy levels (**Fig. 1F, Supplementary Note Figures 2 and 3A**,**B**). Notably, representation of a given cell type in the training data was a major determinant of its prediction accuracy (**Supplementary Note Fig. 3C**), implying higher model performance can be achieved with the incorporation of more datasets.

First we applied the high-hierarchy (i.e. low-resolution, 38 cell types) classifier to our cross-tissue dataset and found high consistency when comparing the predicted cell types to our coarse-grained manual annotations (**Fig. 1G**). Among them, the monocytes and macrophages, which often form a transcriptomic continuum in scRNA-seq datasets due to their functional plasticity, were clearly resolved by CellTypist. Furthermore, as the training dataset of CellTypist contained hematopoietic tissues with definitive annotations for progenitor populations, the classifier was able to resolve the manually annotated progenitors into HSC/MPP, neutrophils and monocytes. Thus CellTypist was able to both reinforce and refine our manual annotations.

To allow for automated annotation of fine-grained immune sub-populations, we next applied the low-hierarchy (high-resolution, 93 cell types and subtypes) classifier, which is able to predict cells into more specific subtypes including subsets of T cells, B cells, ILCs and dendritic cells (**Fig. 1G**). This classification highlighted a high degree of heterogeneity within the T cell compartment, not only distinguishing between αβ and γδ T cells, but also dissecting CD4+ and CD8+ T cell subtypes and their more detailed effector and functional phenotypes. Specifically, the CD4+ T cell cluster was classified as helper, regulatory and cytotoxic subsets, and the CD8+ T cell clusters contained unconventional T cell subpopulations such as NKT. Moreover, CellTypist identified a population of germinal center B cells and also revealed three distinct subsets of dendritic cells - DC1, DC2 and migratory DCs (*21, 22*), again highlighting the granularity CellTypist can achieve.

In summary, we have generated an in-depth map of immune cell populations across human tissues, and developed a framework for automated annotation of immune cell types and subtypes. CellTypist produced expert-grade annotations on our multi-tissue and multi-lineage dataset, and its performance as assessed on multiple metrics was comparable or better relative to other label-transfer methods with minimal computational cost (**Supplementary Note Figures 4**,**5**). This approach allowed us to substantially refine the description of multiple cell subtypes such as the progenitors and dendritic cell subtypes at full transcriptomic breadth, resolving 26 cell states in total across our dataset. This automated annotation forms the basis for further cross-tissue comparisons of cell compartments in the sections below. CellTypist is publicly available on github and pypi, with user-friendly documentation for application to any single cell transcriptomics data sets.

### Tissue-restricted features of mononuclear phagocytes

Mononuclear phagocytes, including monocytes, macrophages and dendritic cells, are critical for immune surveillance and tissue homeostasis. Building on the CellTypist annotation, subclustering of the myeloid subsets revealed further heterogeneity, particularly within macrophages and monocytes (**Fig. 2A-C, fig. S2A**).

**Fig. 2.**
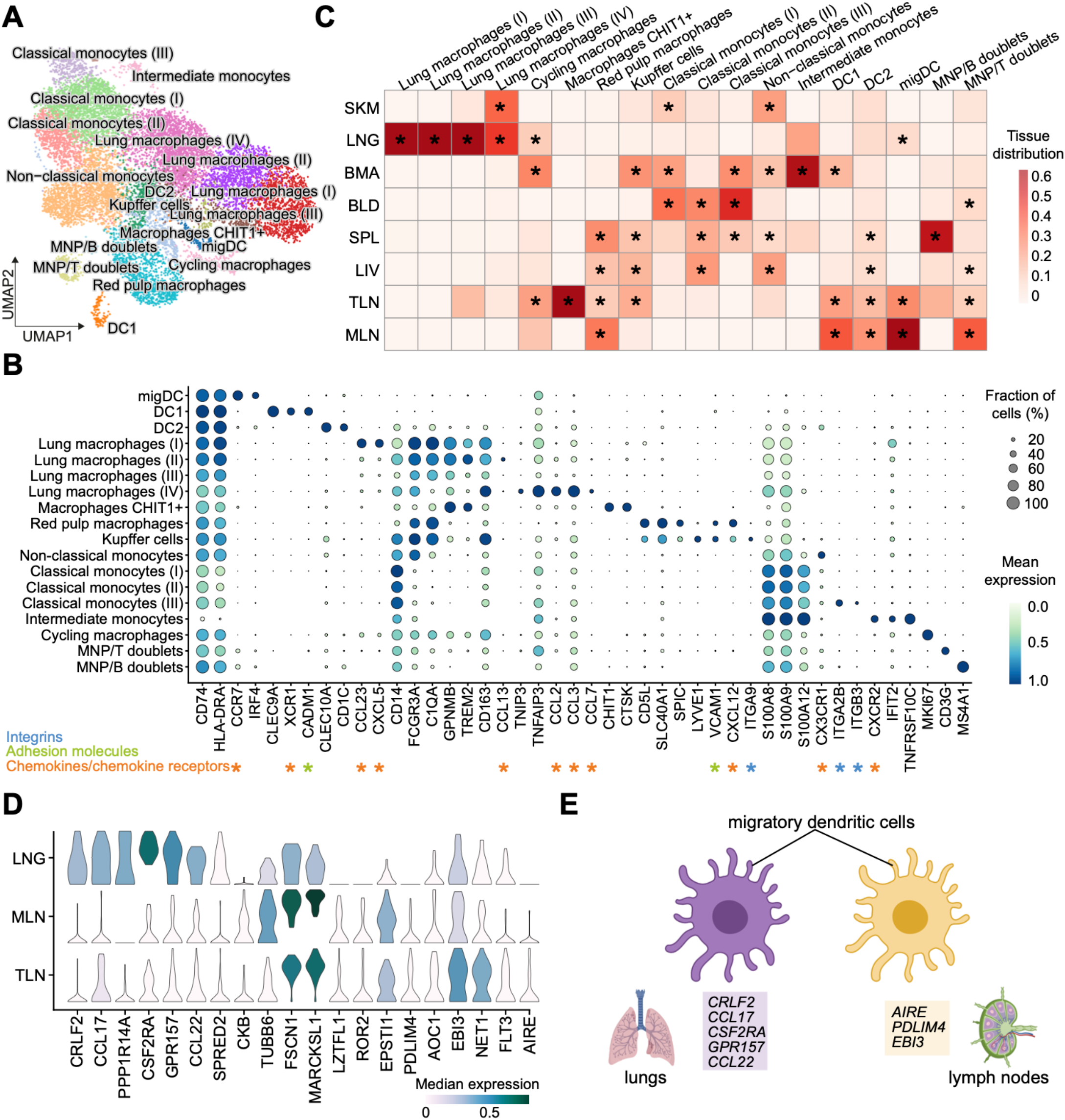
Myeloid compartment across tissues. (**A**) UMAP visualization of the cell populations in the myeloid compartment. (**B**) Dot plot for expression of marker genes of the identified myeloid populations. Color represents maximum-normalized mean expression of cells expressing marker genes, and size represents the percentage of cells expressing these genes. (**C**) Heatmap showing the distribution of each myeloid cell population across different tissues. Cell numbers are normalised with each tissue and later calculated as proportions across tissues. Asterisks mark significant enrichment in a given tissue relative to the remaining tissues (poisson regression stratified by donors, *p* < 0.05). (**D**) Violin plot for genes differentially expressed in migratory dendritic cells across tissues. Color represents maximum-normalized mean expression of cells expressing marker genes. (**E**) Schematic illustration of tissue-specific features of migratory dendritic cells.

Among macrophages, lung-resident cells constituted the majority, and were classified into four major clusters: MAC-I and MAC-II expressing *GPNMB* and *TREM2*, which have been related to alveolar macrophages(*23*) and disease-associated macrophages (*24*), respectively. MAC-III expressing epithelial markers (potentially due to ambient RNA, doublets or the ingestion of epithelial debris), and MAC-IV with unique expression of *TNIP3* (**Fig. 2B**). TNIP3 (TNFAIP3-interacting protein 3) binds to A20, encoded by TNFAIP3, and inhibits TNF, IL-1 and LPS induced NF-kB activation(*25*). Its expression in lung macrophages may be related to underlying pathology as it was primarily detected in donor A29, a multitrauma donor with lung contusions, and bibasal lung consolidation secondary to aspiration of gastric contents (**fig. S2B**). Consistent with this, among lung macrophages, MAC-IV also expressed the highest levels of S100A8/9, which together form calprotectin, an antimicrobial peptide due to its ability to sequester iron and zinc. Notably, MAC-IV also expressed monocyte-recruiting chemokines CCL2 and CCL3 (**Fig 2B**), providing a means of replenishing the lung macrophage pool.

Beyond these lung macrophages, other macrophage subsets in our data also showed a high degree of tissue restriction (**Fig. 2C**). Red pulp macrophages and Kupffer cells mainly populated the spleen and liver, as expected, and shared high expression of *CD5L, SCL40A1* and the transcription factor *SPIC(26)*, with higher expression of *LYVE1* and *ITGA9* distinguishing the Kupffer cells. Notably, a number of macrophages from other tissues, such as the bone marrow and lymph nodes, clustered together with red pulp macrophages and Kupffer cells, pointing to phenotypic similarities across subsets of iron-recycling macrophages (**Fig. 2C**).

We also found a macrophage subtype in the thoracic lymph node which expressed the chitin- and kinin-degrading enzymes *CHIT1* and *CTSK*, respectively (**Fig 2B,C**). Macrophages upregulate *CHIT1* during terminal differentiation, and its expression and activity correlates with lung disease (*27, 28*). The *CHIT1*-expressing macrophages were restricted to the thoracic lymph node (TLN) in our data, while other publications have reported these in the lung (*29*).

Similar to macrophages, analyses of monocytes demonstrated significant heterogeneity. In addition to three subsets of classical *S100A8*/*9*/*12*+ monocytes, we identified one non-classical monocyte subtype expressing *FCGR3A* and *CX3CR1*, and an intermediate monocyte subtype mainly restricted to the bone marrow with modest *CD14* expression (**Fig. 2A,B**). Among dendritic cells, DC1 expressed *XCR1* and *CLEC9A*, consistent with their identity as conventional/classical DCs (cDC1), specialised for cross-presentation of antigen (**Fig. 2B**). Two *CD1C/IRF4+* subsets were evident, DC2 and a *CCR7*+ subset. *CCR7* is upregulated in tissue DCs following TLR or FcγR ligation(*30, 31*), enabling migration towards CCL19/21-expressing lymphatic endothelium and stromal cells in the T cell zone of lymph nodes(*32, 33*). Consistent with this, we observed a marked enrichment of *CCR7*+ migratory DCs (migDCs) in thoracic and mesenteric lymph nodes, with some cells also observed in the lung (**Fig. 2C**). We validated the presence of migDCs in TLN by immunofluorescence (**fig. S2C**).

Interestingly, migDCs showed upregulation of *AIRE, PDLIM4* and *EBI3* in the TLN, and to a lesser extent in the MLN, while in the lung they upregulate *CRLF2* (encoding TLSPR), chemokines (*CCL22, CCL17*), *CSF2RA* and *GPR157* (**Fig. 2D, E**). TLSPR is involved in the induction of Th2 responses in asthma (*34*). Expression of these genes in lung DCs was also observed in our previously published scRNA-seq datasets(*35, 36*) (**fig. S2D**,**E**). These observations suggest that dendritic cell activation coincides with the acquisition of tissue-specific markers that differ depending on the local tissue microenvironment.

Overall, our analysis of the myeloid compartment revealed shared and tissue-restricted features of 18 mononuclear phagocyte cell states, including rare populations of iron-recycling macrophages in mesenteric lymph nodes, chitin-degrading macrophages in thoracic lymph nodes and new subtypes of tissue-specific activated dendritic cells.

### B cell subsets and immunoglobulin repertoires across tissues

B cells, comprising progenitors in the bone marrow, developmental states in lymphoid tissues and terminally differentiated memory and plasma cell states in lymphoid and non-lymphoid tissues, play a central role in humoral immunity *via* the production of antibodies that are tailored to specific body sites. Within CellTypist there are four immature and six mature B cell subsets. Using these models and manual curation, we performed an in depth analysis of the B cell compartment, revealing finer subpopulations across tissues, and describing 12 cell states in total (including two likely physiological doublet B cell states) (**Fig. 3A, fig. S3A**,**B**).

**Fig. 3.**
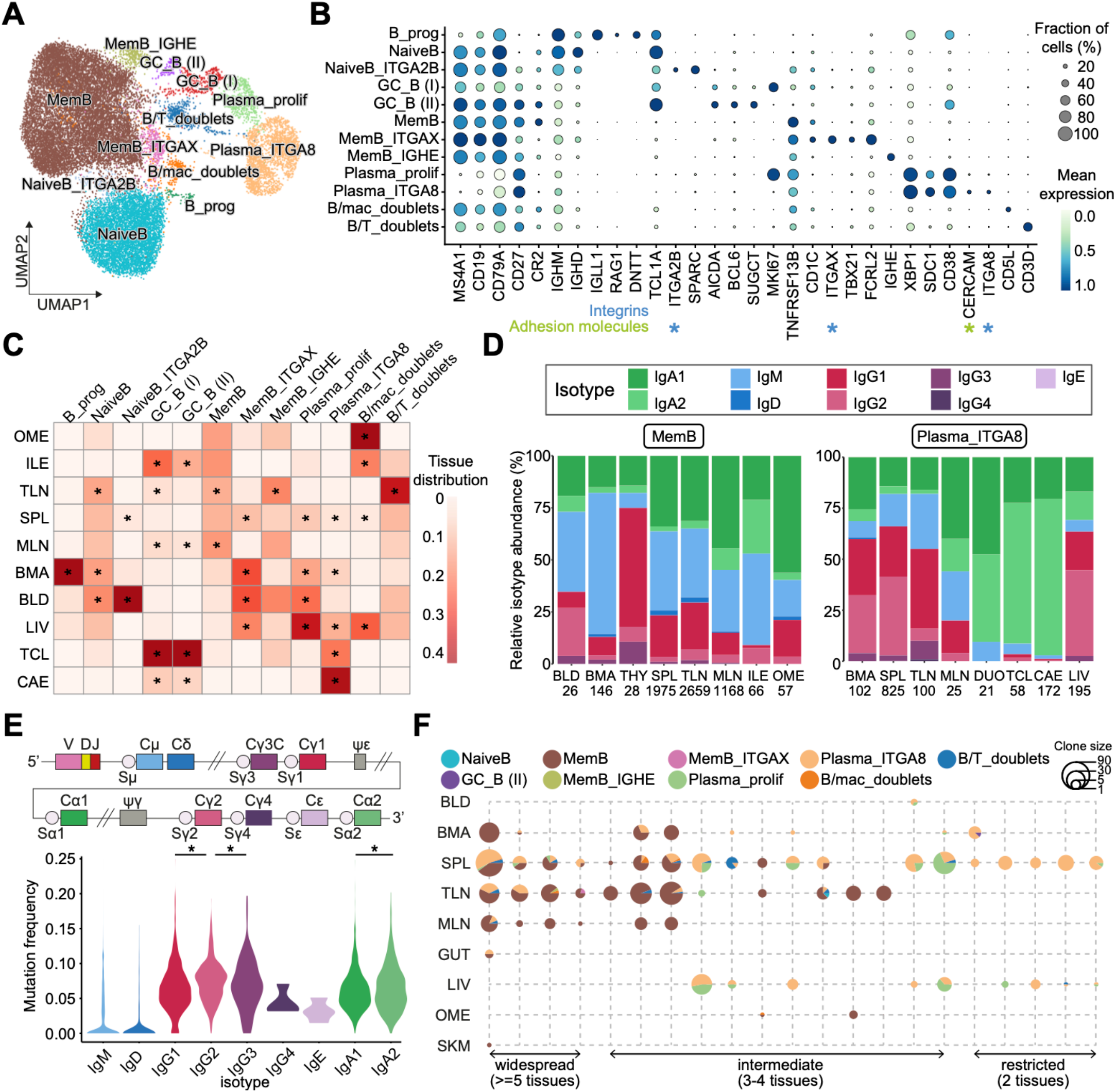
B cell compartment across tissues. (**A**) UMAP visualization of the cell populations in the B cell compartment. (**B**) Dot plot for expression of marker genes of the identified B cell populations. Color represents maximum-normalized mean expression of cells expressing marker genes, and size represents the percentage of cells expressing these genes. (**C**) Heatmap showing the distribution of each B cell population across different tissues. Cell numbers are normalised with each tissue and later calculated as proportions across tissues. Asterisks mark significant enrichment in a given tissue relative to the remaining tissues (poisson regression stratified by donors, *p* < 0.05). (**D**) Stacked bar plots showing the isotype distribution per tissue within the memory B cell cluster and the plasma cell cluster. (**E**) Violin plot of hypermutation frequency on the IgH chain across isotypes. Significant difference between IgG2 and IgG1/3, as well as between IgA2 and IgA1 is marked by asterisks (wilcoxon rank sum test, *p* < 0.05). (**F**) Scatterpie plot showing the tissue distribution and B cell subsets of expanded clonotypes (>10 cells). Each vertical line represents one clonotype. Clonotypes are grouped based on their tissue distributions.

Within the memory B cells, which were characterized by expression of B-cell lineage markers (*MS4A1, CD19*) and *TNFRSF13B*, we found a transcriptomically distinct cluster (MemB_ITGAX) positive for *ITGAX, TBX21* and *FCRL2*, encoding CD11c, T-bet and the Fc receptor-like protein 2, respectively (**Fig. 3B**). CD11c+T-bet+ B cells, also known as age-associated B cells (ABCs), have been reported in autoimmunity and aging (*37*–*39*), and likely correspond to our MemB_ITGAX population. Notably, unlike conventional memory B cells, they showed low expression of *CR2* (encoding CD21) and *CD27* (**Fig. 3B**). Interestingly, this subset was mainly present in the blood, liver and bone marrow, while in secondary lymphoid organs, it was primarily found in the spleen (confirmed by flow cytometry and IF **fig. S3C**,**D**), with only few cells detectable in the thoracic and none in the mesenteric lymph nodes (**Fig. 3C**). This data deepens our understanding of the phenotype of this non-classical subtype of memory B cells, their adhesion properties and tissue distributions.

Within the naive B cell compartment of the blood and spleen, we observed a small population that was distinct from the bulk of naive B cells, with unique expression of *ITGA2B* and *SPARC* (encoding the matricellular protein - secreted protein acidic rich in cysteine - that plays a key role in tissue stroma remodeling) (*40, 41*) (**Fig. 3B**). In addition, we identified two small populations of germinal center B cells expressing *AICDA* and *BCL6* differing in their proliferative states (marked by *MKI67*). Of note, we did not find any differential expression of dark zone and light zone marker genes for these two populations, probably reflecting limited germinal center activity in the adult donors. Moreover, these germinal center populations were present in lymph nodes and diverse gut regions (**Fig. 3C**), presumably representing Peyer’s patches.

We also uncovered two populations of plasmablasts (Plasma_prolif) and plasma cells (Plasma_ITGA8) marked by expression of *CD38, XBP1* and *SDC1*. Whereas the former expressed *MKI67* and were found in the spleen, liver, bone marrow and blood, the latter expressed the integrin alpha-8-encoding gene *ITGA8*, the adhesion molecule *CERCAM* and was detected in the large intestine (CAE and TCL) in addition to the tissues of the Plasma_prolif population (**Fig. 3B,C**). ITGA8+ plasma cells have been recently reported in the context of an analysis of bone marrow plasma cells (*42*), and are likely a long-lived plasma cell population that is resting and tissue-resident. Here we expand their tissue distribution to the liver and gut, and describe their specific clonal distribution pattern below.

B cells have an additional source of variability due to VDJ recombination, somatic hypermutation and class-switching, which can relate to cell phenotype. We performed targeted enrichment and sequencing of BCR transcripts to assess isotypes, hypermutation levels and clonal architecture of the B cell populations described above. This analysis revealed an isotype and subclass usage pattern that related to the cellular phenotypes (**Fig. S4A**). As expected, progenitors and naive B cells mainly showed a subclass preference for IgM and IgD. Interestingly, while memory B cells presented evidence of class switching to IgA1 and IgG1, plasmablasts and plasma cells also showed significant class switching to IgA2 and IgG2.

To determine to what extent this isotype subclass bias correlated with the tissue of origin, we assessed each cell state independently (requiring a minimum cell count of 19 for statistical power). Memory B cells showed a bias towards IgA1 in the omentum and mesenteric lymph node, and towards both IgA1 and IgA2 in the ileum where Peyer’s patches are found (**Fig. 3D**). In the plasma cell compartment, we found an even more striking preference for IgA2 across several gut regions (caecum, duodenum and transverse colon), consistent with the known dominance of this isotype at mucosal surfaces (**Fig. 3D**). IgG-expressing plasma cells were also evident in the large intestine, but not in the duodenum, consistent with previous studies (*15*). Of note, plasma cells in the bone marrow, liver and spleen were composed of over 20% IgG2 subclass. With more limited numbers, we report isotype distributions across tissues for other B cell populations (**Fig. S4B**,**C**): for example, the *ITGAX*+ memory B cells are dominated by IgA subtypes in the thoracic lymph node, and by IgM in the spleen.

Somatic hypermutation (SHM) levels were, as expected, lowest in naive B cells and highest in plasma cells (**Fig. S4D**). Between isotypes and subclasses, SHM did not differ significantly. Nonetheless, there was a tendency towards higher mutation rates in the distal classes IgG2 and IgA2, which are downstream from the IgH locus and can thus accumulate more mutations during sequential switching (*43*) (**Fig. 3E**).

We also explored the occurrence of sequential class switching events in our data by assessing the isotype frequency among expanded clonotypes (>10 cells). Mixed isotype clones are rare in our data, with only a minority detected between IgM and IgD (**Fig. S4E**). We then evaluated the distribution of these expanded clones across tissues and cell types and found three major groups of clones - present in only two tissues, three to four tissues or five or more tissues, respectively (**Fig. 3F**), similar to previously reported patterns of B cell clonal tissue distribution (*44*). While the clones restricted to two tissues, typically between the spleen and the liver or bone marrow, were enriched in plasma cells, those distributed across more than five tissues, including lymph nodes, were over-represented in memory B cells. Together, these findings suggest that tissue-restricted clones could represent a long-term immunological memory maintained by long-lived plasma cells (Plasma_ITGA8 population) resident in the bone marrow and spleen as well as liver in our data.

Overall, we describe 12 cell states in the B cell compartment, with new in depth characterisation of both naive and memory B cell as well as plasma cell subsets. We identify distinct patterns of clone distributions for the more tissue-restricted long-lived Plasma_ITGA8 cells *versus* broad tissue distribution of classical memory B cell clones.

### TCR repertoire analysis reveals patterns of clonal distribution and differentiation across tissues

For annotation of the T cell/innate lymphocyte compartment, as for the B cell compartment, we combined information from CellTypist (27 cell types in this compartment) with in depth manual annotation, which further divided these populations into 15 substates by adding manual inspection to CellTypist scoring (**Fig. 4A**).

**Fig. 4.**
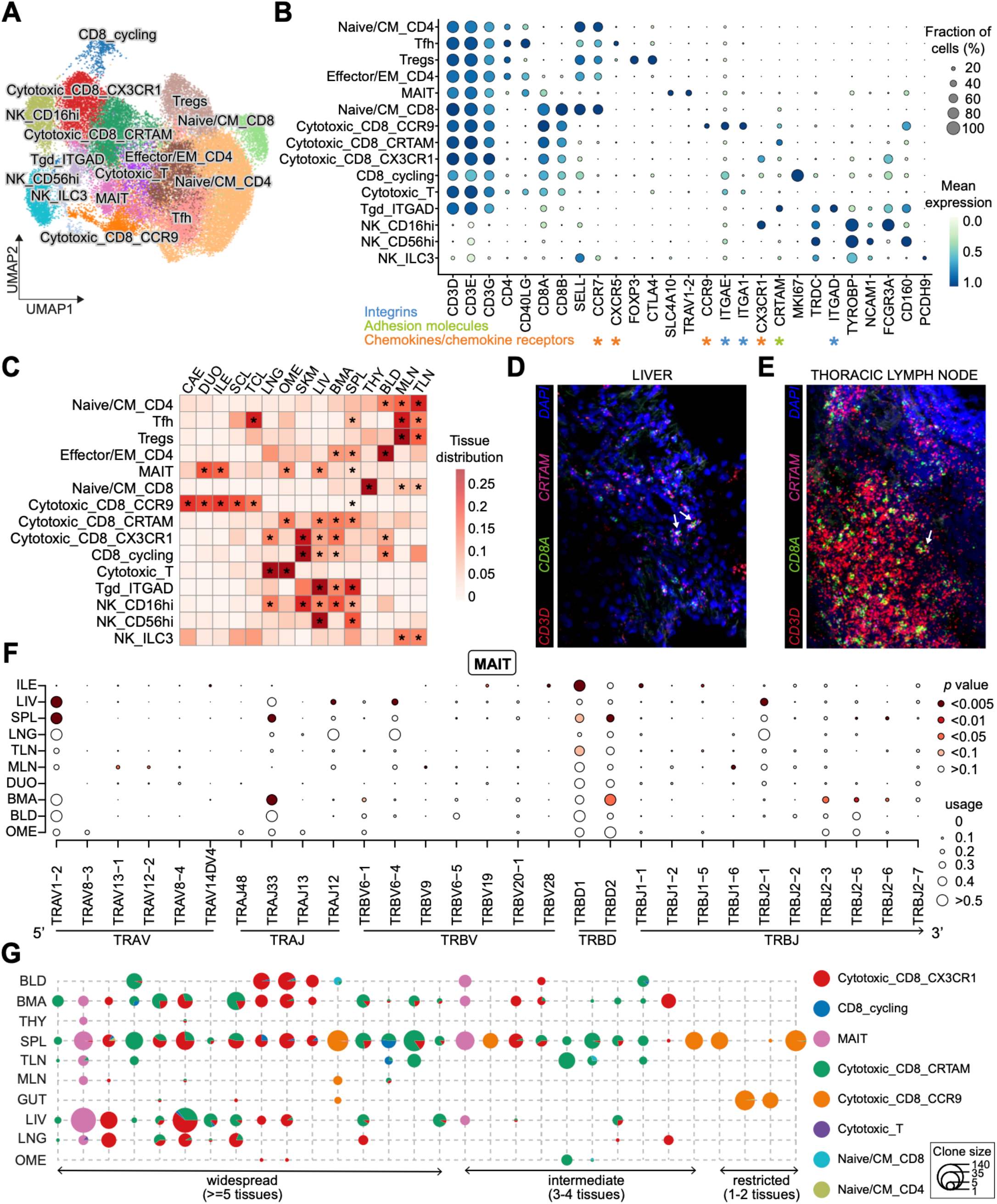
TCR repertoire analysis reveals patterns of clonal distribution and differentiation across tissues. (**A**) UMAP visualization of T cells and innate lymphoid cells (ILCs) across human tissues colored by cell types. (**B**) Dot plot for expression of marker genes of the identified immune populations. Color represents maximum-normalized mean expression of cells expressing marker genes, and size represents the percentage of cells expressing these genes. (**C**) Heatmap showing the distribution of each T cell or ILC population across the different tissues. Cell numbers are normalised with each tissue and later calculated as proportions across tissues. Asterisks mark significant enrichment in a given tissue relative to the remaining tissues (poisson regression stratified by donors, *p* < 0.05). (**D**,**E**) smFISH visualisation of *CD3D, CD8A* and *CRTAM* transcripts, validating this tissue-resident memory CD8+ T cell population in the liver and thoracic lymph node. (**F**) Dot plot denoting the TRA and TRB V(D)J usage across tissues for MAIT cells. Only gene segments with a usage of greater than 10% in at least one tissue are included in the plots, with the sizes of dots indicating the gene segment usage and the colors indicating the significance of the difference between a given tissue and the remaining tissues. Significance is assessed by the generalized linear model stratified by donors with a Poisson structure. (**G**) Scatterpie plot showing the tissue distribution and T cell subsets of expanded clonotypes (>20 cells). Each vertical line represents one clonotype. Clonotypes are grouped based on their tissue distributions.

Naive/central memory CD4+ T cells were transcriptionally close to naive CD8+ T cells as defined by high expression of *CCR7* and *SELL* (**Fig. 4B**). Other CD4+ T cells identified included: follicular helper T cells (Tfh) expressing *CXCR5*, regulatory T cells (Tregs) expressing *FOXP3* and *CTLA4*, and effector memory CD4+ T cells. Within the effector memory CD8+ T compartment, we found three major subsets characterized by expression of the activation molecule *CRTAM* and the chemokine receptors *CCR9* and *CX3CR1*, respectively. The *CCR9+* population, which also expressed tissue-resident markers *ITGAE* and *ITGA1* (**Fig. 4B**), was enriched in the gut regions, whereas the *CX3CR1+* population was ubiquitous (with the exception of lymph nodes) (**Fig. 4C**). A third CD8+ T cell population expressing *CRTAM* was found in both lymphoid and non-lymphoid tissues. We validated and mapped this population of *CD3D+CD8A+CRTAM*+ T cells using single-molecule FISH (smFISH) in the liver (**Fig. 4D**) and thoracic lymph nodes (**Fig. 4E**). Furthermore we validated all three effector memory CD8+ T cell populations at the protein level by flow cytometry of cells purified from human spleen (**fig. S5**). Although we could validate CRTAM at the RNA level by smFISH, the protein could not be detected without stimulation, suggesting that CRTAM is subject to post-translational regulation upon TCR activation. These three distinct cytotoxic effector memory T cell populations may represent different states of tissue adaptation and/or maturation between T effector memory and tissue-resident memory states. We also detected unconventional T cell subsets such as MAIT cells, characterised by expression of *TRAV1-2* and *SLC4A10*, and a γδ T cell population overexpressed the integrin molecule *ITGAD* (CD11d) (**Fig. 4B**). NK cells in our data were represented by two clusters with high expression of either *FCGR3A* (encoding CD16) or *NCAM1* (encoding CD56). We defined the ILC3 population, within a small cluster mixed with NK cells, based on markers including *PCDH9* (**Fig. 4A,B**).

Analyses of tissue distributions of these populations revealed that, whereas the majority of CD4+ T and ILC3 cells were located in the lymph nodes and to some extent in the spleen, cytotoxic T and NK cells were more abundant in the bone marrow, spleen and non-lymphoid tissues (**Fig. 4C**). Notably, the CD4:CD8 ratio in our gut data was lower than previously reported (*45*), which could be related to a lower yield from lamina propria and associated lymphoid tissue due to our pan-tissue dissociation method(*46*).

The analysis of T cell clonal distribution within different tissues of a single individual, and across different individuals is key to understanding T cell-mediated protection. We identified a total of 30,842 cells with productive TCRαβ chains. Chain pairing analysis showed that cells from the T cell clusters mostly contained a single pair of chains (50-60%), with orphan (5-20%) and extra (5-10%) chains present in small fractions of cells (**Fig. S6A**,**B**). Notably, the frequency of extra α chains (extra VJ) was higher than that of β chains (extra VDJ), a phenomenon potentially due to more stringent and multi-layered allelic exclusion mechanisms at the TCRβ locus compared to TCRα (*47*). As expected, the NK and ILC clusters held no productive TCR chains. Within the γδ T cell cluster, only a small proportion had a productive TCR chain, which may result from cytotoxic T cells co-clustering with γδ T cells.

We next examined V(D)J gene usage in relation to T cell identity. In the MAIT population, we detected a significant enrichment of *TRAV1-2*, as expected (**Fig. 4F, Fig. S6C**), and revealed an exciting tissue-specific distribution of *TRAJ33* (spleen) and *TRAJ12* (liver) (**Fig. 4F**). Specifically, V(D)J usage bias analysis across tissues revealed a significant enrichment of *TRAJ12, TRBV6-4* and *TRBJ2-1* in liver MAIT cells *versus TRAJ33* and *TRBJ2-6* in splenic MAIT cells (**Fig. 4F**). This suggests that there may be different antigens for MAIT cells in the spleen *versus* liver corresponding to the different metabolomes in these tissues. In addition, full analysis of the TCR repertoire of the MAIT cells revealed previously unappreciated diversity of V segment usage in the beta chain (**Fig. S6D**).

We then defined clonally related cells based on identical CDR3 nucleotide sequences to investigate their TCR repertoires. Using this approach, we found that clonally expanded cells were primarily from the memory CD8+ T cell compartment, and the MAIT populations mentioned above, with fewer small CD4+ T cell clones (**fig. S6E**). These clonotypes were restricted to single individuals as expected (**fig. S6F**), and within an individual they were distributed across tissues and subsets (**fig. S6G**,**H**). We found a very small number of isolated CD4+ T cell clones that shared Treg and effector T cell phenotypes, illustrating the possibility but low frequency of plasticity or (trans)differentiation from the same naive precursor cell in the periphery (**fig. S6H**).

Focusing on the most expanded clonotypes (>20 cells), the majority were widespread across five or more tissues supporting the systemic nature of tissue-resident immune memory (**Fig. 4G**). Moreover, we found that several *CCR9*+ CD8+ T cell-enriched clonotypes were shared across different gut regions, and that several clonotypes present across both the liver and lung consisted of a mixture of cells of the *CRTAM*+ and *CX3CR1*+ CD8+ T cell populations. This suggests that a single naive CD8+ T cell precursor can give rise to diverse cytotoxic T cell states, which harbour immune memory across multiple non-lymphoid tissues, again emphasizing plasticity of phenotype and location within a clone.

In summary, we describe 15 T/innate cell states in our data by integrating three sources of information: CellTypist logistic regression models, V(D)J sequencing data and manual inspection. This yields new insights into the MAIT cell compartment and its antigen receptor repertoire distribution that is markedly different in spleen *versus* liver. For CD4+ T cells, we find rare sharing of Tregs and effector memory cells within the same clone. We show that for the cytotoxic T cell memory compartment there is broad sharing of clones across gut regions for *CCR9*+ CD8+ T cell-enriched clonotypes, and mixed *CRTAM*+ and *CX3CR1*+ CD8+ T cell clonotypes with broad tissue distributions.

### Two distinct subsets of γδ T cells across human tissues

γδ T cells are at the interface between adaptive and innate immune functions, and have recently attracted attention due to their potential for cell-based therapies. Accurate identification of γδ T cells in single-cell transcriptomics studies remains challenging due to their transcriptional similarity to cytotoxic αβ T cells and NK cells. Here we manually annotated a γδ T cell cluster expressing *ITGAD* which encodes CD11d, part of a CD11d/CD18 heterodimer with the potential to interact with the cell adhesion molecule VCAM1 (*48*). In the locations where γδ T cells were identified (liver, spleen and bone marrow), resident macrophages, such as red pulp macrophages, expressed high levels of *VCAM1* (**Fig. 2B, Fig. 5A**), indicating a potential interaction between these cells. Due to the lack of a robust commercially available anti-CD11d antibody, we validated this population by flow sorting and qPCR on sorted CD3+TCRγδ+ and CD3+TCRαβ+ cells from cryopreserved spleen samples from three donors (**Fig.S7**). As a small fraction of αβ T cells, marked by low expression of CD52 and CD127, were noted to express *ITGAD*, the CD3+TCRαβ population was split into CD52-CD127- and CD52+CD127+ subpopulations. In keeping with our scRNAseq data, *ITGAD* expression was high in CD3+TCRγδ and CD52-CD127-CD3+TCRαβ (**Fig. 5B**), providing additional evidence for the specific expression of this integrin alpha subunit in this subpopulation of γδ T cells. In order to further elucidate the identity of this interesting lymphocyte compartment, we carried out γδ TCR sequencing in selected spleen, bone marrow and liver samples. The γδ TCR sequencing data was subjected to a customized analysis pipeline we developed and optimised on the basis of the use of Cell Ranger followed by contig re-annotation with dandelion (see Methods), facilitating the full recovery of γδ chains in our data. This analysis confirmed that the majority of productive γδ TCR chains originated from the *ITGAD-*expressing γδ T cells (**Fig. 5C, fig. S8A**), further supporting the robust identification of this population. Overall, these multiple data support a γδ T cell population in the liver, spleen and bone marrow that may play a role in tissue homeostasis *via* interaction with tissue-resident macrophage populations.

**Fig. 5.**
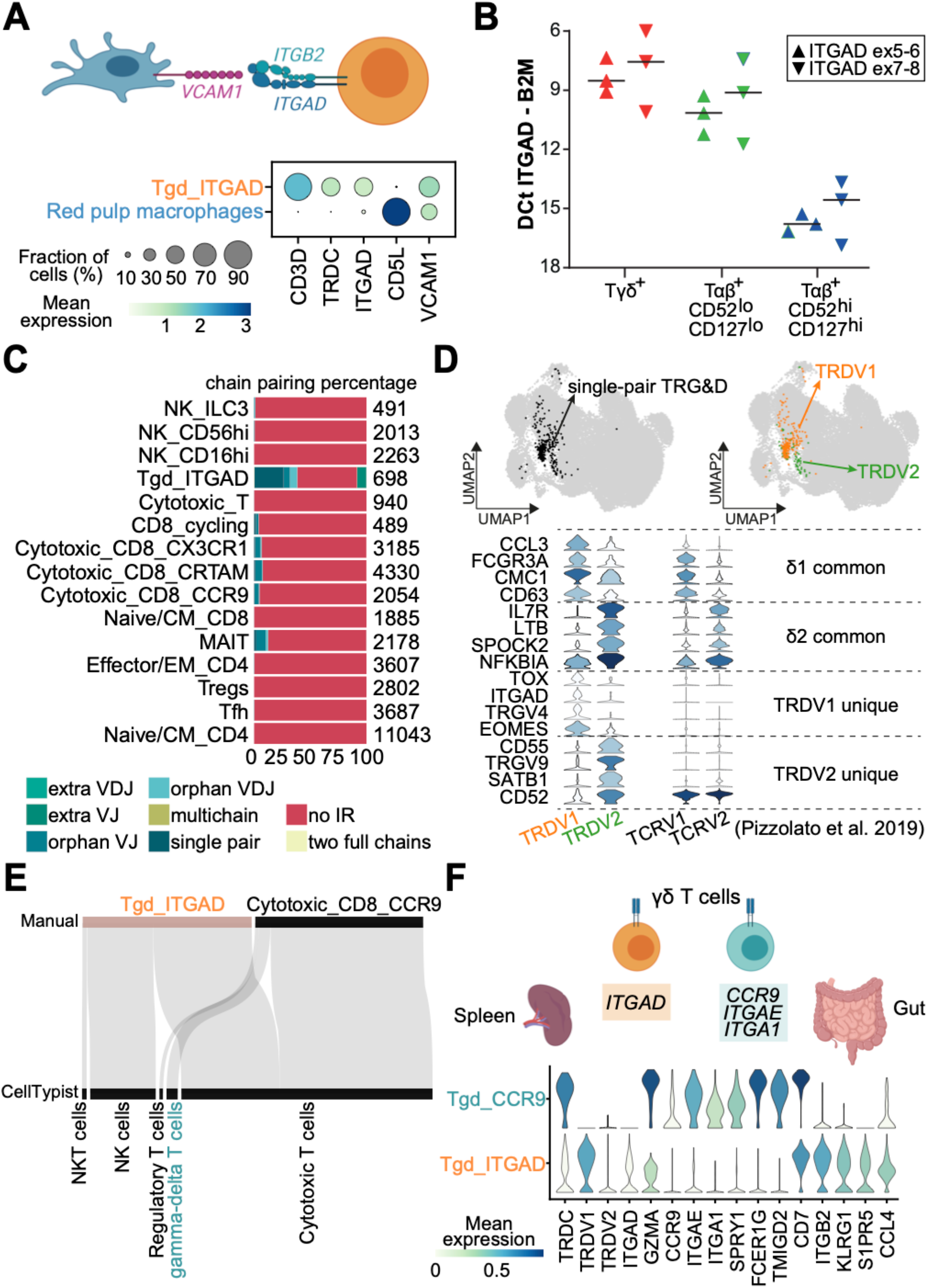
Two distinct subsets of γδ T cells across human tissues. (**A**) Dot plot showing the expression of *VCAM1* by *CD5L+* macrophages (marking red pulp macrophages) and *TRDC* and *ITGAD* transcripts by γδ T cells in the spleen. (**B**) qPCR validation of *ITGAD* in T cell populations as measured by two assays to *ITGAD* and normalised to the housekeeping gene *B2M*. (**C**) Stacked bar plot depicting the distributions of γδ TCR chain pairing types across cell populations. No IR = No immune receptor. (**D**) UMAP plots showing the cells with single-pair γδ TCR chains (upper left) and the subdivision of these cells into the TRDV1 and TRDV2 clusters (upper right). Bottom violin plots demonstrate the expression of common and unique marker genes of the two clusters as compared to the γδ T subsets from Pizzolato et al. (**E**) Sankey plot of the *CCR9*-expressing γδ T cells identified using CellTypist. (**F**) Stacked violin plot for expression of marker genes of the two identified γδ T cell subsets. Color represents maximum-normalized mean expression of cells expressing marker genes.

γδ T cells in the blood have been shown to comprise two transcriptomically distinct subsets according to their T cell receptor V gene: TCRVδ1 or TCRVδ2(*49*). Our unique data allowed us to explore in detail the characteristics of these two subsets. Specifically, analyses of γδ TCR chains revealed a further subdivision of cells with productive single-pair γδ chains into the TRDV1 and TRDV2 subclusters, consistent with their separation trend in the UMAP representation (**Fig. 5D**). Comparisons of the two subpopulations with the blood TCRVδ1 and TCRVδ2 showed that they shared common markers for TCRVδ1 (*CCL3, FCGR3A, CMC1, CD63*) and TCRVδ2 (*IL7R, LTB, SPOCK2, NFKBIA*). Moreover, in comparison to blood, γδ T cells derived from the spleen, liver and bone marrow were uniquely overrepresented by specific markers, such as expression of *TOX* and *TRGV4* in the TRDV1 subcluster, as well as *CD55* and *TRGV9* in the TRDV2 subcluster (**Fig. 5D**).

In addition to *ITGAD-*expressing γδ T cells, CellTypist predicted a different group of cells (sitting within the *CCR9*+ CD8+ T cell cluster) as γδ T cells (**Fig. 5E, fig. S8B**). This CellTypist-annotated γδ T cell population was primarily derived from the gut and had high expression of the gut homing receptor CCR9, as well as several integrin-related genes including *ITGAE* and *ITGA1* (**Fig. 5F**).

Altogether, dissection of these different tissue-resident γδ T cell populations in human tissues for the first time showcases how a combination of CellTypist-based automated annotation, expert-driven cluster analysis and TCR sequencing can synergize to dissect specific and functionally relevant aspects of cell identities.

## Discussion

Here, we present the first multi-donor/multi-tissue study of immune cells across the human body. By sampling multiple organs from the same individuals, which allows for robust control of age, gender, medical history, drug exposure and sampling backgrounds, we have been able to reveal novel tissue-specific expression patterns across the myeloid and lymphoid compartments.

Accurately assigning detailed cell identities in scRNA-seq experiments is a major challenge. To overcome this, we have developed CellTypist, a framework for automated immune cell type annotation. In addition to identifying major cell types, CellTypist is able to perform fine-grained cell subtype annotation that is typically very time-consuming and requires expert knowledge. This has largely been made possible due to the curation of a large collection of studies across a range of tissues with in-depth immune cell analysis comprising 93 harmonized cell type labels. Currently, manual curation still has a role to play in revealing specific cell subtypes that may be absent from the database/training set, and we showcase such an example in the γδ T cell compartment. To address this, in the longer term, the CellTypist models will be periodically updated and extended to include further immune and non-immune sub-populations, as more data become available.

Using CellTypist combined with manual annotation, we have dissected the transcriptomic features of 49 immune cell types and states across tissues. Within the myeloid compartment, macrophages showed most prominent features of tissue specificificity. Red pulp macrophages and Kupffer cells, shared features related to iron-recycling(*50*) with macrophages in other locations such as the bone marrow and mesenteric lymph nodes, suggesting that macrophages participate in iron metabolism across a wide range of tissues. Among lung macrophages, we found a subtype with high TNIP3 and calprotectin expression. Elevated serum calprotectin has been proposed as a biomarker of bacterial pneumonia(*51*), predicts pulmonary exacerbations and lung function decline in cystic fibrosis(*52*), and an S100A8/9 high macrophage subset has been identified in COVID19 bronchiolar lavage fluid(*53*). Lastly, we characterized tissue-specific features of migratory dendritic cells (CCR7+) revealing specific expression of *CRLF2, CSF2RA* and *GPR157* in the lung, and expression of *AIRE* in the thoracic lymph nodes. These novel migratory dendritic cell states are interesting targets for future in depth functional characterisation in the context of allergy, asthma and other related pathologies (*54, 55*).

In the lymphoid compartment, we combined single-cell transcriptome and VDJ analysis, which allowed the phenotypic dissection of adaptive immune cells using complementary layers of single cell genomics data. Of note, we detected a subset of memory B cells expressing *ITGAX* (CD11c) and *TBX21* (T-bet) that resemble age-associated B cells (ABCs) previously reported to be expanded in ageing(*38*), following malaria vaccination (*56*) and in systemic lupus erythematosus (SLE) patients (*57*). In our data, these B cells did not show clonal expansion, suggesting that they may be present at low levels in healthy tissues and expand upon challenge as well as ageing. BCR analysis revealed isotype usage biased towards IgA2 in gut plasma cells, which may be related to structural differences (*58*) or higher resistance to microbial degradation as compared to IgA1 (*59*). Cross-tissue clonal distribution of B cells and plasma cells has been previously shown to follow two different patterns (*44*). In our study, we primarily detected shared clones between the spleen and lymphoid organs but were also able to detect a degree of sharing with non-lymphoid tissues including the gut. Notably, when we incorporated cell identities into the clonotype analysis, we observed that broadly distributed clones were geared towards the B cell memory phenotype while tissue-restricted clones were mainly the plasma cells. The plasma cells are therefore likely long-lived plasma cells contributing to immune memory *via* more restricted tissue localisation as compared to memory B cells.

In the T-cell compartment, cross-tissue and cross-cell type TCR sharing revealed previously unappreciated insights into the plasticity and distribution of T cell subtypes. Sharing between *CRTAM*+ and *CX3CR1*+ subtypes of CD8+ T cells supports the possibility that these transcriptomically distinct populations may represent different stages of migration or tissue adaptation, such as the differentiation of effector memory T cells (TEM) to resident memory T cells (TRM) (*60*). Lastly, using a combination of automated annotation, manual curation and γδ TCR sequencing, we found several distinct TRDV1 and TRDV2 subsets of γδ T cells showing distinct integrin gene expression and tissue distributions.

In summary, this study reveals previously unrecognized features of tissue-specific immunity in the myeloid and lymphoid compartments, and provides a comprehensive framework for future cross-tissue cell type analysis. Further investigation of human tissue-resident immunity is needed to determine the effect of important covariates such as underlying critical illness, donor age and gender as well as considering the immune cell activation status, to gain a defining picture of how human biology influences immune functions. Applications of a deep characterisation of the features of immune cells across human tissues include implications for engineering of cells for therapeutic purposes and addressing cells to the intended tissue locations, for understanding tissue-specific features of infection as well as the distinct modes of vaccine delivery to tissues.

## Acknowledgements

We thank the deceased organ donors, donor families and the extended Cambridge Biorepository for Translational Medicine team for access to the tissue samples.

We thank the CellTypist Annotation Team (composed by Simone Webb, Laura Jardine (Muzlifah Haniffa laboratory), Benjamin Stewart, Kelvin Tuong (Menna Clatworthy laboratory), Regina Hoo (Roser Vento-Tormo laboratory), Kylie James, Jongeun Park, Elo Madissoon, Waradon Sungnak, Rasa Elmentaite, Peng He (Sarah Teichmann laboratory) for contributions to the harmonization of CellTypist labels. We acknowledge J. Eliasova and BioRender for graphical images. We thank Carlos Talavera-López and Natsuhiko Kumasaka for discussions on data analysis. We also acknowledge the support received from the Cellular Generation and Phenotyping (CGaP) core facility, the Cellular Genetics Informatics team and Core DNA Pipelines at the Wellcome Sanger Institute and the Cambridge NIHR BRC Cell Phenotyping Hub (Dept. Medicine, University of Cambridge). We are grateful to the CZI Immune Ageing Seed Network consortium for helpful discussions.

## Supplementary Materials

Materials and Methods

Supplementary Figures

Supplemental Note on CellTypist

## Materials and methods

### Tissue acquisition

All work was completed under ethically approved studies. Tissue was obtained from deceased organ donors following circulatory death (DCD) via the Cambridge Biorepository for Translational Medicine (CBTM, https://www.cbtm.group.cam.ac.uk/), REC 15/EE/0152. Briefly, donors proceeded to organ donation after cessation of circulation. Organs were then perfused *in situ* with cold organ preservation solution and cooled with topical application of ice. Samples for the study were obtained within 60 minutes of cessation of circulation and placed in University of Wisconsin (UW) organ preservation solution for transport at 4°C to the laboratory. Gut samples were taken from the locations indicated in Figure 1. Additional samples were obtained from the left lower lobe of the lung and the right lobe of the liver. Skeletal muscle was taken from the third intercostal space and bone marrow was obtained from the vertebral bodies. In addition, two donor-matched blood samples were taken just prior to treatment withdrawal, under REC approval 97/290.

### Donor metadata table

| Donor ID | Sex | Age | Primary cause of death | Multitrauma | Days in hospital | BMI | CMV/EBV/TOXO | Smoking | Alcohol (u/day) | Vasoactive agents | Antibiotics within 2 weeks of death | Steroids |
| --- | --- | --- | --- | --- | --- | --- | --- | --- | --- | --- | --- | --- |
| A29 | F | 65-70 | ICH | Y | 2 | 30-35 | CMV+/EBV+/TOXO- | N/K | <1 | Desmopressin<br>Metaraminol | Nil | N |
| A31 | M | 50-55 | ICH | Y | 8 | 30-35 | CMV+/EBV+/TOXO- | Y | <1 | Noradrenaline | Co-amoxiclav,<br>Tazocin | N |
| A35 | M | 60-65 | ICH | N | 2 | 20-25 | CMV-/EBV+/TOXO- | Y | >9 | Noradrenaline<br>Desmopressin | Gentamicin,<br>Flucloxacillin | Y<br>Dexamethasone |
| A37 | F | 55-60 | ICH | N | 3 | 20-25 | CMV-/EBV+/TOXO- | Y | >9 | Noradrenaline<br>Metaraminol | Co-amoxiclav | N |
| A36 | M | 70-75 | ICH | N | 5 | 25-30 | CMV-/EBV+/TOXO+ | Y | <2 | Noradrenaline<br>Metaraminol | Amoxicillin (pre-admission),<br>Flucloxacillin,<br>Gentamicin,<br>Clarithromycin,<br>Co-amoxiclav | Y<br>Prednisolone pre-treatment |
| A52 | M | 60-65 | ICH | N | 2 | 35-40 | CMV-/EBV+/TOXO- | N/K | N/K | Noradrenaline<br>Metaraminol<br>Desmopressin | Co-amoxiclav | N |

F = Female; M = Male; ICH = intracranial haemorrhage; CMV = Cytomegalovirus; EBV = Epstein-Barr virus; TOXO = Toxoplasmosis; N/K=not known; Y = Yes; N = No;

### Tissue processing

All tissues were processed using a uniform protocol. Briefly, solid tissues were transferred to a 100mm tissue culture dish, cut into small pieces and transferred to C-tubes (Miltenyi Biotec) at a maximum of 5g/tube in 5mL of X-vivo15 media containing 0.13U/m Liberase TL (Roche), 10U/mL DNase (benzonase nuclease, Millipore/Merck) supplemented with 2% (v/v) heat-inactivated fetal bovine serum (FBS; Gibco), penicillin (100 U/ml, Sigma-Aldrich), streptomycin (0.1 mg/ml, Sigma-Aldrich), and 10mM HEPES (Sigma Aldrich). The samples were then dissociated using a GentleMACS Octo dissociator (Miltenyi Biotec) using a protocol that provided gradual ramping up of homogenisation speed along with 2x 15 minute heating/mixing steps at 37°C. Digested tissue was filtered through a 70-μm MACS Smartstrainer (miltenyi biotec) and washed with media containing 2mM EDTA, prior to washing with PBS. A ficoll density centrifugation step (400g for 30min at RT) was performed to isolate mononuclear cells (MNCs). After gradient centrifugation, cells were washed once with PBS prior to counting and resuspending in 10X buffer (PBS containing 0.04% (v/v) BSA).

Bone marrow aspirates and blood samples were diluted 1:1 with PBS and layered directly onto ficoll for mononuclear cell isolation as described above. Cells taken from the interphase layer were washed with PBS and exposed to the same enzymatic conditions as solid tissues by resuspending cell pellets in tissue dissociation media containing liberase TL for 30 minutes at 37°C prior to counting and resuspending in 10X buffer.

### Single-cell RNA sequencing

For scRNA-seq experiments, single cells were loaded onto the channels of a Chromium chip (10x Genomics) for a target recovery of 5,000 cells. Single-cell cDNA synthesis, amplification, and sequencing libraries were generated using the Single Cell 5′ Reagent Kit following the manufacturer’s instructions. The libraries from up to eight loaded channels were multiplexed together and sequenced on an Illumina NovaSeq. The libraries were distributed over eight lanes per flow cell and sequenced at a target depth of 50,000 reads per cell using the following parameters: Read1: 26 cycles, i7: 8 cycles, i5: 0 cycles; Read2: 98 cycles to generate 75-bp paired-end reads. VDJ libraries for B and T cells were sequenced on HiSeq 4000.

### Single-cell RNA sequencing data pre-processing

scRNA-seq data was aligned and quantified using the cellranger software (version 3.0.2, 10x Genomics Inc.) using the GRCh38 human reference genome (official Cell Ranger reference, version 1.2.0). Cells with fewer than 1,000 UMI counts and 600 detected genes were excluded from downstream analysis. scTCR-seq data was aligned and quantified using the cellranger-vdj software (version 2.1.1, 10x Genomics Inc). scBCR-seq data was aligned and quantified using the cellranger-vdj software (version 4.0, 10x Genomics Inc). For TCRγδ we implemented a customized pipeline due to the cell ranger vdj annotation algorithm being tuned towards alpha/beta TCR chains. Briefly, TCR gamma/delta libraries were mapped with cell ranger 4.0.0, using the 10x VDJ 4.0.0 reference. All contigs deemed high quality were selected, and reannotated with IgBlast via the workflow provided in dandelion 0.1.3 (https://github.com/zktuong/dandelion). We provide an example notebook showcasing this workflow (https://sc-dandelion.readthedocs.io/en/latest/notebooks/gamma_delta.html).

### Doublet detection

Doublet detection was performed on a per sample basis using the scrublet algorithm (https://github.com/AllonKleinLab/scrublet,(*61*)) with percolation as previously described (*62*). Briefly, scrublet scores were obtained per cell and the percolation step was performed on over-clustered data using the *scanpy*.*tl*.*louvain* function from the scanpy package. Each cluster was subsequently separately clustered again, yielding an over-clustered manifold, and each of the resulting clusters had its Scrublet scores replaced by the median of the observed values. The resulting scores were assessed for statistical significance, with P values computed using a right-tailed test from a normal distribution centred on the score median and a median absolute deviation (MAD)-derived standard deviation estimate. The P-values were corrected for false discovery rate with the Bonferroni procedure, and a significance threshold of 0.01 was imposed. Cells with a Bonferroni-corrected P-value under 0.01 were excluded from downstream analysis.

### Clustering, batch alignment and annotation

Downstream analysis included data normalisation (*scanpy*.*pp*.*normalize_per_cell* method, scaling factor 10000), log-transformation (*scanpy*.*pp*.*log1p*), variable gene detection (*sc*.*pp*.*highly_variable_genes*), data feature scaling (*scanpy*.*pp*.*scale*), PCA analysis (*scanpy*.*pp*.*pca*, from variable genes), and Leiden graph-based clustering (*scanpy*.*tl*.*leiden*, clustering resolution manually adjusted) performed using the python package scanpy (version 1.6.0). Data integration across donors was done using batch-balanced KNN (https://github.com/Teichlab/bbknn). Cluster cell identity was assigned by manual annotation using known marker genes as well as computed differentially expressed genes (DEGs). Differential expression across clusters was assessed using rank biserial correlation (https://github.com/Teichlab/rbcde) and *markers*.*py* functions from the thymus atlas (https://github.com/Teichlab/thymusatlas) (*14*). Cross-tissue differential expression was assessed using a ridge regression model that included an interaction term for tissue and cell type. For each tissue, genes were ranked according to their coefficients specific to the given interaction term associated with the tissue of interest. To achieve a high-resolution annotation, we sub-clustered T, B and myeloid cells and repeated the procedure of variable gene selection, which allowed for fine-grained cell type annotation. Donor-dependent batch effects were aligned using the *scanpy*.*pp*.*bbknn* function and we used the batch-aligned manifold to annotate cell types.

### CellTypist

Full details on the data collection, processing, curation, model training, and testing of the CellTypist pipeline can be found in the **Supplementary Note**. Briefly, immune cells from 20 tissues of 19 studies were collected and harmonised into consistent labels with the inputs from experts followed by semi-supervised label propagation to non-annotated cells. These cells were randomly split into equal-sized mini-batches, with each batch containing 1,000 cells. In each epoch, 100 such batches were sequentially trained by the l2 regularized logistic regression using stochastic gradient descent learning. This step was repeated 30 epochs, enabling the CellTypist models to see cell numbers with six orders of magnitude. Lastly, feature selection was performed to choose the top 500 genes from each cell type by ranking the genes according to their absolute weights associated with the given cell type, and the union of these genes across cell types were supplied as the input for a second round of training, with the detailed training procedure being constant as the initial round of training.

### scTCR-seq downstream analysis

VDJ sequence information was extracted from the output file “filtered_contig_annotations.csv” using the scirpy package(*63*). We determined productive TCR chain pairing features using the *scirpy*.*tl*.*chain_pairing()* function and selected cells with a single pair of productive αβ TCR chains for downstream analysis. Bias in VDJ usage across tissues was estimated using the generalized linear model (glm) on the basis of the Poisson family. Specifically, for each gene segment, we calculated the number of cells with this segment in each donor of a given tissue, and then compared their distributions with the remaining tissues which were stratified by donors as well. During the glm fitting, the total number of cells in each donor of a given tissue was logarithmized and used as the offset, accounting for the variance imposed by rate estimation (glm(counts∼tissues + offset(log(total_counts)), family = ’poisson’) in R). Clonotypes were determined using the *scirpy*.*pp*.*tcr_neighbors()* function using the CDR3 nucleotide sequence identity from both TCR chains as a metric.

### scBCR-seq downstream analysis

VDJ sequence information was extracted from the output file “filtered_contig_annotations.csv”. Further single-cell VDJ analysis for B cells was performed broadly as described previously (*15, 64*), with all sequences from a given patient grouped together for analysis. AssignGenes.py(*65*) and IgBLAST(*66*) were used to reannotate IgH sequences prior to correction of ambiguous V gene assignments using TIgGER (v1.0.0)(*67*). Clonally-related IgH sequences were identified using DefineClones.py with a nearest neighbour distance threshold of 0.15 before running CreateGermlines.py (ChangeO)(*68*) to infer germline sequences for each clonal family and calculate somatic hypermutation frequencies with observedMutations (Shazam)(*68*). IgH diversity analyses were performed using the rarefyDiversity and testDiversity of Alakazam (v1.0.2; (*68*)). scVDJ sequences were then integrated with single-cell gene expression objects by determining the number of high quality annotated IgH, IgK or IgL per unique cell barcode. If more than one contig per chain was identified, metadata for that cell was ascribed as “Multi”. To assess clonal relationships between scRNA-seq clusters, co-occurrence of expanded clone members between cell types and tissues was reported as a binary event for each clone that contained a member within two different cell types or tissues in single-cell repertoires.

### Single molecule FISH

Samples were either snap frozen in chilled isopentane (−40°C for striated muscle, -70°C for other tissues) or fixed in 10% NBF, dehydrated through an ethanol series, and embedded in paraffin wax. Samples were run using the RNAscope 2.5 LS fluorescent multiplex assay (automated). Briefly, FFPE tissue sections (5 μm) and fresh frozen tissue sections (10um) were cut. Fresh frozen tissues were pre-treated offline (4% PFA fixation 4°C 15 mins followed by 90mins at room temperature, sequential dehydration steps (50%, 70%, 100%, 100% ethanol, air dry)) and protease III was used. FFPE tissues required no pretreatment offline, but a Heat Induced Epitope Retrieval (HIER) step was performed by the instrument for 15mins using Epitope Retrieval 2 (ER2) at 95°C. These tissues also had protease III treatment. RNAscope probes used included Hs-CD3D-C2 (599398-C2), Hs-CD8A-C3 (560398-C3), Hs-CRTAM (430248). Opal fluorophores (Opal 520, Opal 570 and Opal 650) were used at 1:300 dilution. Slides were imaged on the Perkin Elmer Opera Phenix High-Content Screening System, in confocal mode with 1 μm z-step size, using 20X (NA 0.16, 0.299 μm/pixel) and 40X (NA 1.1, 0.149 μm/pixel) water-immersion objectives.

### Flow cytometry

Spleen, bone marrow and thoracic lymph nodes from additional donors different to the scRNA-seq study were used to validate the discovered cell populations. The mononuclear cells (MNCs) were either stained ex vivo or post activation with PMA+I (eBioscience, Cell Stimulation Cocktail) for two hours. Cells were washed with PBS and then stained with the live/dead marker Zombie Aqua for 10 minutes at room temperature, and then washed with PBS+0.5%FCS. The MNCs were stained in PBS+0.5% FCS at 4°C for 45 minutes with the following panels of antibodies:

CD8 panel: CD3-BUV395 (SK7, BioLegend), CD56-BUV737 (NCAM16.2, BD Horizon), CCR9-BV421 (L053E8, BioLegend), CD4-BV605 (OKT4, BioLegend), TCRgd-Fitc (B1.1, Invitrogen), CX3CR1-PE (2A9-1, BioLegend), CRTAM-PECy7 (CR24.1, Invitrogen), CD16-APC (3G8, BioLegend), CD8-APCCy7 (RPA-T8, BioLegend).

B cell panel: IgD-BUV395 (IA6-2, BD Horizon), CCR7-BV421 (G043H7, BioLegend), CD3-BV605 (SK7, BioLegend), CD11c-BV785 (3.9, BioLegend), CD27-PE (0323, eBioscience), CD19-APC (HIB19, BioLegend), Tbet-PECy7 (eBio4810, eBioscience,Tbet staining was done after the surface staining using the eBioscience Foxp3 transcription factor staining buffer kit).

Cells were fixed with PBS+0.25%PFA and stored at 4°C until they were run on the Fortessa flow cytometry instrument, located in the Cambridge NIHR BRC Cell Phenotyping Hub. Spleen MNCs were used for single stain controls to calculate compensation and FMOs were used to calculate background fluorescence. FlowJo was used to analyse the flow cytometry data.

### qPCR

qPCR was performed to validate the existence ITGAD-expressing γδ T cells in the spleen using three additional samples that were different to those used in the scRNA-seq study. Spleen MNCs were stained with the live/dead marker Zombie Aqua for 10 minutes at room temperature, and then washed with PBS+0.5%FCS. Cells were then stained with the following antibodies at 4°C for 45 minutes: CD56-BV421 (HCD56, BioLegend), CD4-BV605 (OKT4, BioLegend), TCRgd-Fitc (B1.1, Invitrogen), TCRab-PerCPCy5.5 (IP26, BioLegend), CD52-PE (MHCD5204, Life Technologies), CD127-PECy7 (eBioRDR5, eBioscience), CD8-APC (RPA-T8, BioLegend), CD3-APCfire (UCHT1, BioLegend). Cells were washed with PBS+0.5% FCS and passed through a Celltrics (Partec) 30 μm filter prior to cell sorting. Cell sorting was performed on a BD Fusion 4 laser sorter and an example of the gating strategy used is shown in (**fig. S7**).

Sorted cells were pelleted at 600g for 5 minutes and lysed in RNA lysis buffer and frozen until RNA could be extracted. RNA was extracted using a Zymo Research RNA micro kit with on column DNAse digestion. The standard protocol was followed and RNA eluted in 11 μl of water. This RNA was then used to make cDNA using both oligo dT and random hexamer primers with the reverse transcriptase SuperscriptIII. Probes to B2M (housekeeping gene) and two assays to ITGAD were purchased from ThermoFisher, and qPCR reactions were performed in duplicate with the following recipe: 8 μl master mix, 0.7 μl probe, 4.3 μl water and 3 μl cDNA. A ThermoFIsher QuantStudio7 instrument was used for the qPCR, and the Ct values were determined with the DCt being calculated as the Ct of ITGAD - Ct of B2M.

### Immunofluorescence

Spleen and thoracic lymph node samples from unrelated donors were fixed in 1% paraformaldehyde (Electron Microscopy Services) for 24 hours followed by 8 hours in 30% sucrose in PBS. 30µm sections were permeabilized and blocked in 0.1M TRIS, containing 0.1% Triton (Sigma), 1% normal mouse serum, 1% normal rat serum and 1% BSA (R and D). Samples were stained for 2h at RT in a wet chamber with the appropriate antibodies, washed 3 times in PBS and mounted in Fluoromount-G® (Southern Biotech). Images were acquired using a TCS SP8 (Leica, Milton Keynes, UK) confocal microscope. Raw imaging data were processed using Imaris (Bitplane).

Antibodies used: CD3-AF488, clone UCHT1, 1/100 dilution, BioLegend; CD1c-PE, clone L161, 1/50 dilution, BioLegend; CCR7-PE, clone 3D12, 1/50 dilution, eBioscience; CD19-AF594, clone HIB19, 1/100 dilution, BioLegend; CD11c-APC, clone MJ4-27G12, 1/100 dilution, Miltenyi; HLA-DR-AF647, clone TAL 1B5, 1/100 dilution, abcam. Nuclei were stained with Hoechst 33258, 1/10,000 dilution (Biotum).

