## Supplementary Note for "Cross-tissue immune cell analysis reveals tissue-specific adaptations and clonal architecture in humans"

#### Supplementary Note on CellTypist:

##### Introduction

With the growing size of single-cell RNA sequencing (scRNA-seq) datasets and their wide applications in tissue and disease biology (*69*, *70*), fast and accurate cell type annotation becomes of crucial value in order to accelerate the interpretation of newly generated scRNA-seq datasets. Various approaches have been put forward to perform the matching of cellular identities between datasets (*71*). Other efforts have focused on building scRNA-seq references for cell type classification using neural networks (*71*, *72*), such as a recent approach which integrated query datasets with the reference atlases using conditional neural network models (*73*).

In this study, we focused on immune cells and their large variety of subtypes. Immune cells are ubiquitous and mobile across tissues, with specific adaptations to corresponding local environments. This leads to a high degree of cell type heterogeneity, which is further augmented by other factors such as developmental lineage dynamics. Despite this heterogeneity, immune cells can still be grouped into cell types characterised by expression of definitive markers, functional roles, and parent lineages. Therefore, in order to achieve an accurate and specific cell type identification, domain-specific knowledge is always necessary.

To annotate cell types with expert-level details, we introduce CellTypist, a lightweight and directly interpretable pipeline for automatic annotation of scRNA-seq data on the basis of logistic regression classifiers optimised by the stochastic gradient descent (SGD) algorithm. CellTypist currently includes a wide assortment of immune cell types collected from 20 tissues across 19 studies. Extensive model tuning and optimisation is performed to ensure its applicability, with the derived model easily updatable for further releases by incorporating new cell annotations, as well as by including non-annotated cells which in future iterations may be described as novel or specific cell types. Notably, our current CellTypist release involves both low- and high-resolution models which classify cells with coarse and fine granularities, respectively. CellTypist can be readily installed from *https://pypi.org/project/celltypist*.

##### Methods

##### Dataset compilation and label standardisation

scRNA-seq data were collected from 19 publications covering 20 different tissues (**Supplementary Note Figure 1A**). A raw count matrix was obtained for each dataset and subsequently combined across datasets based on their common genes.

In order to focus the model's training data on *bona fide* immune cells, the combined expression matrix was filtered to include only cells expressing *PTPRC*, a general marker for immune cells, as well as not expressing *EPCAM* and *PDGFRA*, markers for epithelial cells and fibroblasts, respectively. In addition, for the datasets which were already annotated in the original publications, only cells identified as immune cell types were included. Exceptions to these rules were "Epithelial cells", "Endothelial cells" and "Fibroblasts" which were retained in the reference dataset to serve as umbrella categories representing fall-backs for non-immune cell types.

For each of these datasets, meta-information was also collected, including the tissues of origin and cell type annotations where possible. In order to train CellTypist models using uniform cell type labels, cell identities across datasets were summarised into consistent names with the knowledge inputs from experts (**Supplementary Note Figure 1B**). These labels encompass two levels of hierarchies: a high-hierarchy (low-resolution) level which includes a total of 38 broad cell types; and a low-hierarchy (high-resolution) level which comprises 93 detailed cell types and subtypes through subdivision of broad cell types. These two levels are arranged hierarchically, such that the low-hierarchy annotations are able to consistently match corresponding high-level classes (**Supplementary Note Table 1**).

##### Propagating annotations to non-annotated cells

After label standardisation, a small subset of cells included in CellTypist still did not obtain cell type labels. These datasets were also subject to the same expression filtering as in "***Dataset compilation and label standardisation***". Given that the non-annotated cells may contain similar cell types as those in the annotated cells, we next sought to minimise the label duplication, facilitating the incorporation of both known and yet-to-be-annotated cell identities into the CellTypist models (**Supplementary Note Figure 1B**). Specifically, the non-annotated cells from each combination of tissue and dataset were clustered independently using a canonical Scanpy pipeline. The resulting clusters were then compared with their predicted cell type labels which were inferred from the CellTypist models trained from the annotated cells (for details of model training, see the next section of "***Model training***"). For a given cluster where at least 75% of its cells matched a specific low-hierarchy annotation label, the whole cluster was annotated as such, and meanwhile was assigned a corresponding high-hierarchy cell type label. For the remaining clusters where this condition was not met, we assigned them cell type labels at the high-hierarchy level where possible, through the same procedure as the low-hierarchy labels. This resulted in a final set of harmonised labels between unannotated and annotated cells for a total of 738,647 cells, leaving only six unknown clusters totalling 2,590 cells with undetermined identities.

##### Model training

Different classifiers for cell type predictions have been described (*71*, *74*). Of note, high performances can be achieved even when the classifiers are constructed using canonical machine learning methods, notably the logistic regression models (*75*, *76*). We based the models of CellTypist on a logistic regression framework with several adaptations.

First, randomly sampled mini-batches, instead of the whole training dataset, were used during the training procedure. This approach not only bypassed the possible memory excess when modelling our large dataset, but also ensured the fast convergence not readily available for datasets with hundreds of thousands of cells. Each mini-batch comprised 1,000 cells sampled from the whole dataset, and in a single epoch 100 mutually exclusive mini-batches were sequentially trained. This step was repeated 30 epochs, enabling the CellTypist models to see cell numbers with six orders of magnitude. In practice, the number of epochs needed will be fewer, with the performance plateau reached within 10 epochs (1,000 iterations) (**Figure 1F**), again highlighting the usefulness of the mini-batch training approach in CellTypist.

Second, SGD algorithm was used in combination with the mini-batch training to derive the solutions of the model cost/loss function. This was implemented using the scikit-learn package in Python (*77*) by the "*partial_fit*" method from the class "*SGDClassifier*". SGD also allows for online training, meaning that if new data with no novel labels are feeded, it can be easily incorporated into the model.

Third, L2 regularization was imposed on the logistic models to make the predictions more applicable to external test datasets. This also allows each gene in the model to have a weight of greater than 0 such that more genes can be utilized when predicting test data with varying numbers of input features. The regularization term (alpha) was chosen by training the models with alpha set to 0.01, 0.001, 0.0001, 0.00001 or 0.000001, and the alpha yielding the best performance on an independently left-out data (10% of the total dataset) was chosen as the optimal hyper-parameter. Ultimately, the alpha was set to 0.0001 for the low-hierarchy model and 0.001 for the high-hierarchy model.

Last, feature selection was conducted before the final models were trained. Specifically, we performed an initial training based on the entire gene set, and selected the top 500 genes from each class (cell type) by ranking the genes according to their absolute weights associated with the given class. After combining the genes from all the cell types, a total of 7,560 genes were obtained and later supplied as the input to a second round of training. This step effectively reduces the complexity of the sample space and emphasises the dominant contributions of highly informative genes to the classification of cell identities.

Before model training, the dataset was normalised to 10,000 counts per cell and log-transformed (with a pseudocount of 1). To enable the fast convergence using the optimal learning rate, as well as to ensure a comparable scale of weights across genes when L2 regularization was applied, expression of each gene was standardised to a mean of zero and unit variance. Furthermore, the mean and standard deviation of all genes are recorded and will be applied to the shared genes between the model and query data in the prediction step.

**Cell type prediction**

Before the prediction, the input test data was normalised to 10,000 counts per cell and log-transformed (with a pseudocount of 1). Only genes shared between the CellTypist model and the input data were used in the downstream prediction. For each gene, as noted in "***Model training***", we standardised it by subtracting the mean and scaling the standard deviation using the corresponding mean and standard deviation recorded in the training step for that gene. For each cell type involved in the model, the decision scores of the test cells are defined as the linear combination of the scaled gene expression and the model coefficients associated with the given cell type ("*decision_function*" from the class "*SGDClassifier*" in sklearn), and the probabilities are calculated by transforming the decision scores with a sigmoid function ("*scipy.special.expit*" in scipy). The two metrics are recorded in CellTypist outputs. Finally, the cell type with the maximal decision score (or probability) is selected as the predicted identity for the query cell. Of note, for each cell, we didn't sum up the probabilities to one across cell types to provide a quantification of the confidence score of each cell type, enabling the examination of novel cell types in the test data.

**Over-clustering and majority voting**

The prediction step is performed to infer the identities of input cells, which renders the prediction of each cell independent. To combine the cell type predictions with the cell-cell transcriptomic relationships, CellTypist offers a majority voting approach based on the idea that transcriptionally similar cells in the query dataset are more likely to form a (sub)cluster regardless of their individual prediction outcomes. In this study, the query data was first over-clustered using the Leiden algorithm on the basis of an existing neighborhood graph in the input object ("*scanpy.tl.leiden*" in Scanpy) with the resolution set to 20. If no neighborhood graph exists for the input data (such as the input of a count matrix), a neighborhood graph will be constructed before the over-clustering ("*scanpy.pp.neighbors*" in Scanpy). Each resulting subcluster was then assigned the identity supported by the dominant cell type predicted for this subcluster. Through this step, distinguishable small subclusters will be assigned distinct cell type labels, and homogenous subclusters will be assigned the same labels and iteratively converge to a bigger cluster.

**Benchmarking with other label-transferring methods**

We focused on the comparisons among five methods: CellTypist, traditional logistic regression (lr) classifier, support vector machine (svm) classifier, Azimuth (*78*), and scNym (*79*). To this end, 10,000 cells were randomly sampled from our compiled dataset as an independent test dataset. We further generated three training datasets with the sizes being 5,000, 50,000, and 250,000 cells respectively, through sampling the cells from the remaining dataset. This allows us to examine the effect of sizes of training datasets on the prediction accuracy, representing small, medium and big training datasets, respectively.

To make the comparisons unbiased across different methods, both the training and test data were properly preprocessed beforehand (the time used for preprocessing is not included in the benchmarking of running time): i) For CellTypist, lr and svm, the training data was normalised and scaled as in "***Model training***". The test data was normalised in the same way while scaled using the recorded mean and standard deviation as in "***Cell type prediction***"; ii) For scNym, the training and test datasets were both normalised to 1,000,000 counts per cell as suggested by the scNym guidelines and then log-transformed (with a pseudocount of 1); iii) For Azimuth, the training and test datasets were both normalised to 10,000 counts per cell and log-transformed (with a pseudocount of 1). For all the five methods, we used the same set of highly variable genes extracted from the reference object ("*scanpy.pp.highly_variable_genes*" in Scanpy).

We split the whole label-transferring procedure into the "training" and "prediction" steps. Moreover, we define a "user time" as the time needed for a user to get their prediction results after supplying the test data to the programs. This is critical as the user time is more related with the user experience in practice. **Supplementary Note Figure 5A** lists the detailed splitting for the five methods. Specifically, in CellTypist, lr and svm, the training steps are only dependent on the training data, while in scNym and Azimuth, the training steps rely on both the training and test data ("*scnym_api(task='train')*" in scNym, and "*FindTransferAnchors*" in Seurat, respectively). Therefore, from the perspective of a user, the user time in CellTypist, lr and svm equals to the prediction time while to the sum of training and prediction time in scNym and Azimuth.

For each method, we recorded both the training and prediction time, as well as the predicted cell types for the test data. The performance was then assessed for each cell type separately using three metrics: precision ("*sklearn.metrics.precision_score*"), recall ("*sklearn.metrics.recall_score*"), and F1 score ("*sklearn.metrics.f1_score*").

#### Results

##### CellTypist performance on a fully annotated mouse reference

To test the standalone applicability of CellTypist in a setting with diverse known cell types, we trained a model based on the *Tabula Muris (80)* dataset. Prior to combining the droplet-based and plate-based datasets, we removed any cell types represented by fewer than ten cells. We then used 90% of this combined dataset to train a model as described in "***Model training***", and used the remaining 10% to test the model performance. We observed that the model's accuracy measured by the F1 score increased to approximately 95% only after 50 iterations (**Supplementary Note Figure 2A**), and most cell types showed F1 scores of greater than 0.75 (**Supplementary Note Figure 2B**). Importantly, we found that the obtained F1 score for the test dataset was correlated with the cell type size (number of cells within a given cell type), with cell types represented by at least 100 cells having scores of greater than 0.75 (**Supplementary Note Figure 2B**). These results showed that CellTypist is capable of deriving a high-accuracy model using a training dataset with highly diverse cell types.

##### CellTypist performance on the cross-tissue immune reference

We next examined the performance of CellTypist on our assembled immune cell atlas. For an independently left-out dataset (10%), the CellTypist models trained from the remaining dataset (90%) demonstrated the precision of 0.96 and 0.9 at the high- and low-hierarchy levels, respectively

(**Supplementary Note Figure 3A**). The recall scores were relatively lower, but still reached 0.87 and 0.84 at the two levels respectively (**Supplementary Note Figure 3B**). Further summarising the two metrics into the F1 score, the CellTypist models overall exhibited the F1 scores of 0.94 and 0.88 at the two levels (**Figure 1F**). Examination of the F1 score for each cell type annotated in the models revealed that part of the models' prediction errors came from a low number of cells associated with certain labels (**Supplementary Note Figure 3C**), indicating a future need of collecting more rare cell types.

Benchmarking with other label-transferring methods reveals that when the data size is small (5,000 cells), CellTypist has a comparable performance as compared to the traditional logistic regression, Azimuth and scNym, all of which outperform svm (**Supplementary Note Figure 4**). When the data size is medium (50,000 cells) or large (250,000 cells), CellTypist has a similar performance with the traditional logistic regression and scNym, which is slightly better than Azimuth and much better than svm. Importantly, our mini-batch training approach with SGD learning dramatically decreases the time needed for the model training and thus represents a more scalable method for large-scale scRNA-seq datasets (**Supplementary Note Figure 5B**). In terms of the user time (defined in "***Benchmarking with other label-transferring methods***"), as with canonical machine learning methods, CellTypist predicts the test data much more efficiently and quickly than Azimuth and scNym (see the user time marked by asterisks in the **Supplementary Note Figure 5B**), largely due to the independence between the data training and prediction steps in CellTypist.

#### Discussion

Automatic cell type prediction of unknown cells is among the most important approaches to making full use of the hard-earned knowledge from existing single-cell transcriptomics datasets, in particular when studying cells which are as diverse as those from the immune system (*81*).

The application of CellTypist to the *Tabula Muris* data highlights the advantages of training a model using datasets from different tissues and generation protocols. We expect the continuous inclusion of new datasets in our reference data will further refine the prediction models, as well as our own notions of cell type standards. We also expect future releases of CellTypist models will converge on more accurate solutions by increasing the number of cells supporting a given cell type. Meanwhile, encompassing more tissues and cell types will improve the models' ability to generalise to new datasets and to delineate the gene expression profile defining each cell type included. Maintaining this will require continuous curation by specialised researchers as well as the research community.

Our strategy for cell type classification bears some limitations. While our models are readily interpretable and can achieve a low computational footprint during data training and prediction, more advanced algorithms may improve the prediction accuracy particularly for lowly represented cell types, as well as incorporation of a hierarchical classification structure within the same models. The use of the mini-batch strategy may also be updated in future releases when more data are included which may necessitate an unbiased representation of cell types during sampling.

A

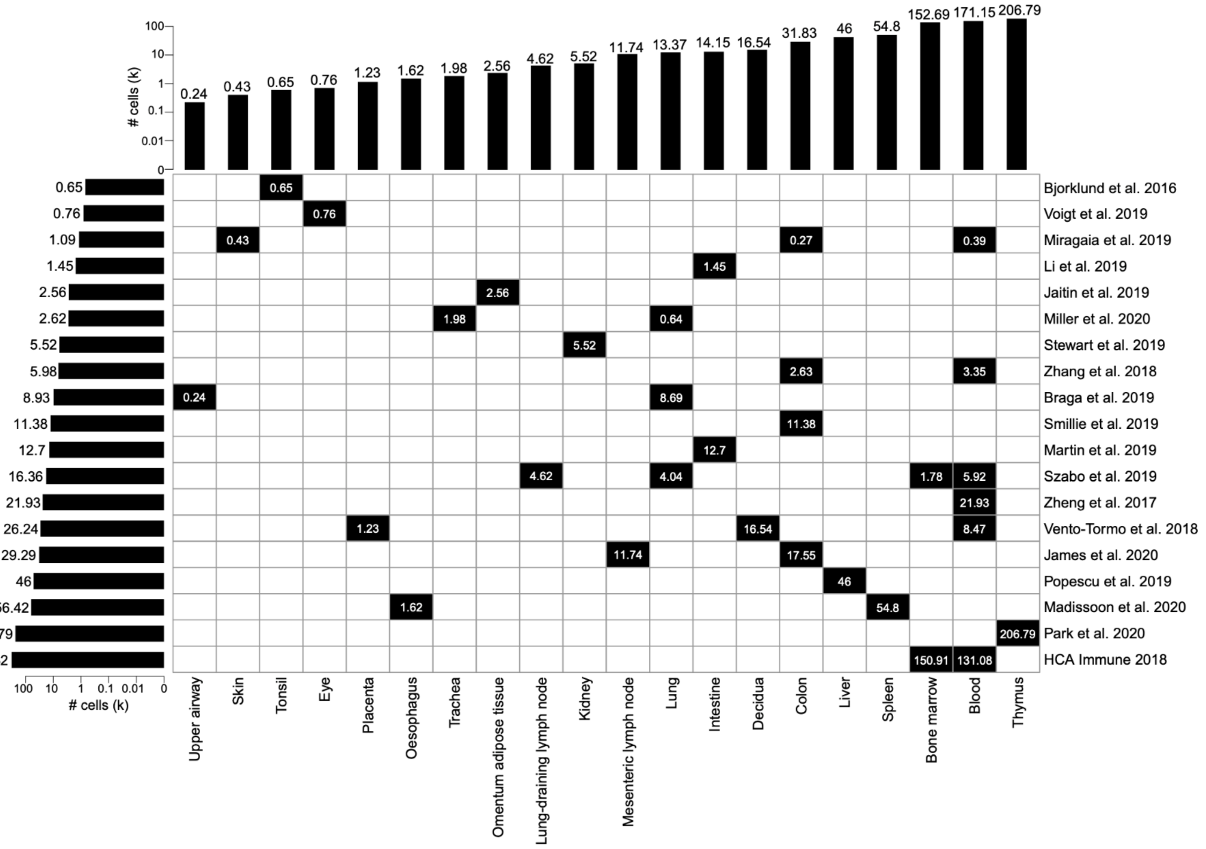

### cells (k)
0.24
0.43
0.65
0.76
1.23
1.62
1.98
2.56
4.62
5.52
11.74
13.37
14.15
16.54
31.83
46
54.8
152.69
171.15
206.79
Bjorklund et al. 2016
Voigt et al. 2019
Miragaia et al. 2019
Li et al. 2019
Jaitin et al. 2019
Miller et al. 2020
Stewart et al. 2019
Zhang et al. 2018
Braga et al. 2019
Smillie et al. 2019
Martin et al. 2019
Szabo et al. 2019
Zheng et al. 2017
Vento-Tormo et al. 2018
James et al. 2020
Popescu et al. 2019
Madissoon et al. 2020
Park et al. 2020
HCA Immune 2018
Upper airway
Skin
Tonsil
Eye
Placenta
Oesophagus
Trachea
Omentum adipose tissue
Lung-draining lymph node
Kidney
Mesenteric lymph node
Lung
Intestine
Decidua
Colon
Liver
Spleen
Bone marrow
Blood
Thymus


B

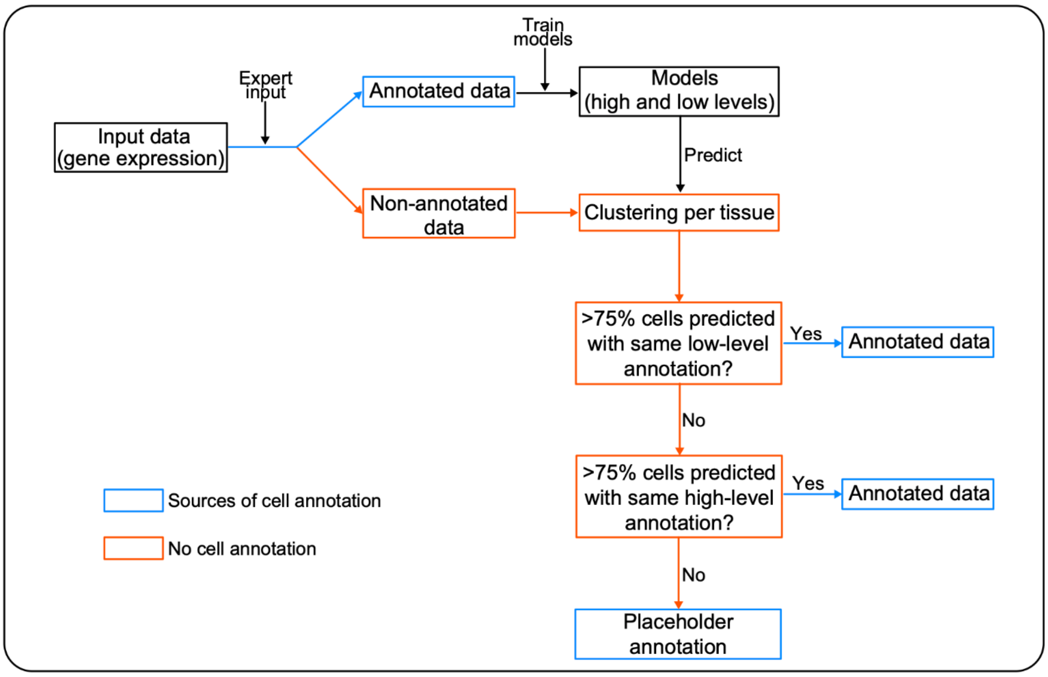

Input data (gene expression)
Expert input
Annotated data
Non-annotated data
Train models
Models (high and low levels)
Predict
Clustering per tissue
>75% cells predicted with same low-level annotation?
Yes
Annotated data
No
>75% cells predicted with same high-level annotation?
Yes
Annotated data
No
Placeholder annotation
Sources of cell annotation
No cell annotation

**Supplementary Note Figure 1**. **Summary of the assembled datasets and cell type label standardisation.** (**A**) Heat map showing the number of cells in each combination of dataset (rows) and tissue (columns), as well as the total number in each dataset (horizontal bar plot) and each tissue (vertical bar plot). Cell numbers are denoted in units of thousands. (**B**) Schematic of the CellTypist pipeline to harmonise cell type labels across datasets including the propagation of known cell type labels to non-annotated cells.

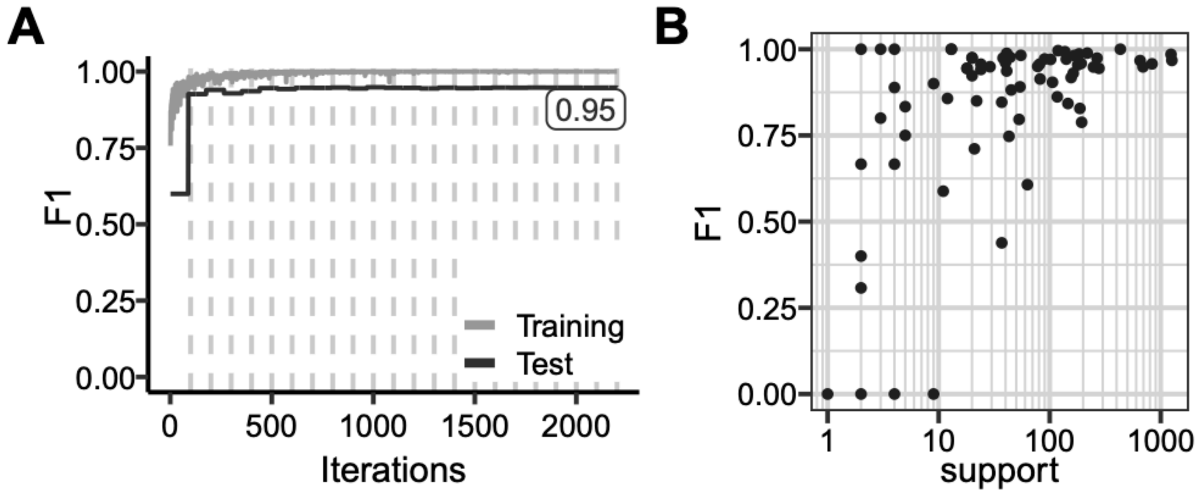
A
B
F1
1.00
0.75
0.50
0.25
0.00
0
500
1000
1500
2000
Iterations
0.95
Training
Test
1
10
100
1000
support

**Supplementary Note Figure 2**. **Evaluation of CellTypist by predicting cell identities for a highly diverse *Tabula Muris* reference.** (**A**) Testing accuracy of a model measured by F1 score based on 90% of the total *Tabula Muris* dataset during training. The number marks the final obtained accuracy. (**B**) F1-score for each tested cell type as a function of its representation in the whole *Tabula Muris* dataset (corresponding to 10% of the total).

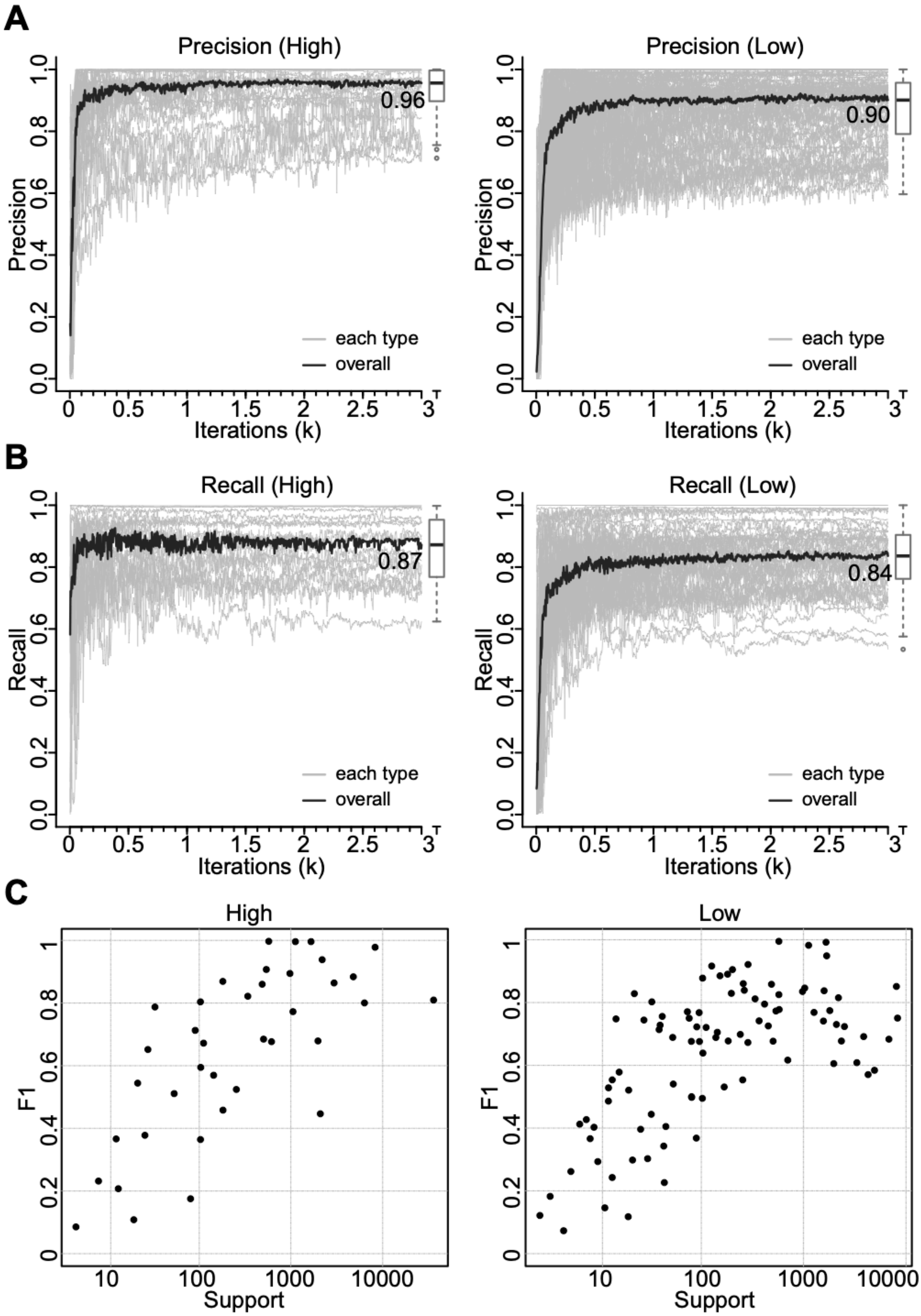
A
Precision (High)
Precision
Iterations (k)
0.96
each type
overall
Precision (Low)
0.90
B
Recall (High)
Recall
0.87
Recall (Low)
0.84
C
High
Low
F1
Support

**Supplementary Note Figure 3**. **Building a human immune reference to predict immune cell identities.** (**A,B**) Performance curves showing the precision (**A**) and recall (**B**) scores at each iteration of training using mini-batch stochastic gradient descent for high- and low-hierarchy CellTypist models, respectively. The black curves represent the median scores averaged across the individual scores of all predicted cell types (grey curves). (**C**) F1-score for each tested high-hierarchy (left) or low-hierarchy (right) cell type as a function of its representation in the compiled human immune datasets test set (corresponding to 10% of the total).

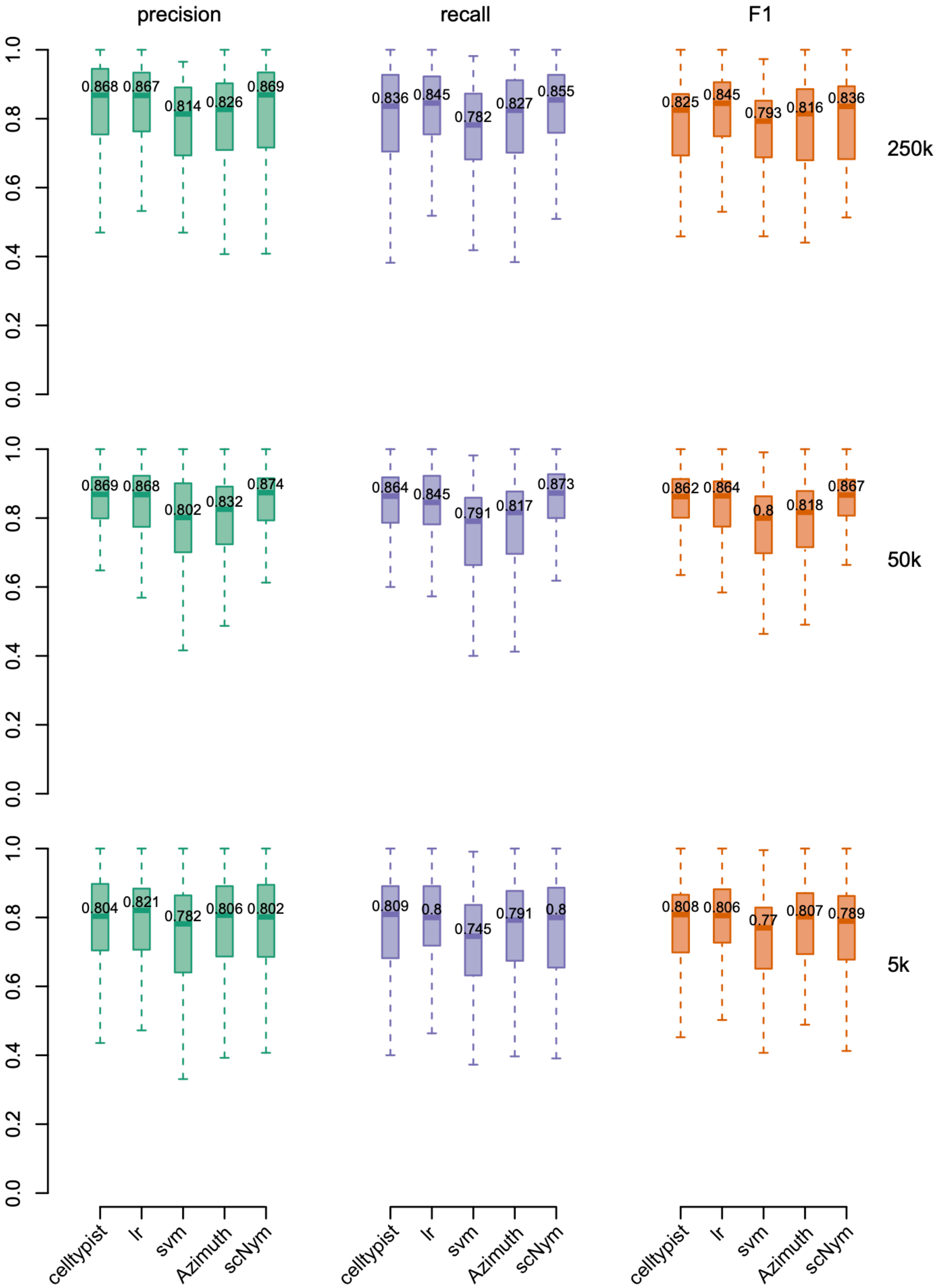

precision
recall
F1
250k
50k
5k
0.868
0.867
0.814
0.826
0.869
0.836
0.845
0.782
0.827
0.855
0.825
0.845
0.793
0.816
0.836
0.869
0.868
0.802
0.832
0.874
0.864
0.845
0.791
0.817
0.873
0.862
0.864
0.8
0.818
0.867
0.804
0.821
0.782
0.806
0.802
0.809
0.8
0.745
0.791
0.8
0.808
0.806
0.77
0.807
0.789
celltypist
lr
svm
Azimuth
scNym

**Supplementary Note Figure 4**. **Benchmarking of CellTypist accuracy with other methods.** Box plots showing the prediction precision (left), recall (center) and F1 score (right) for the training dataset with 5,000 (lower), 50,000 (middle), and 250,000 (upper) cells, respectively. Five methods are assessed and the median value of these metrics across individual cell types is shown for each method.

**A**

| | CellTypist | lr | svm | Azimuth | scNym |
|---|---|---|---|---|---|
| **Training** | *celltypist.train* | *LogisticRegression.fit* | *LinearSVC.fit* | *FindTransferAnchors* | *scnym_api(task='train')* |
| **Prediction** | *celltypist.annotate* | *LogisticRegression.predict* | *LinearSVC.predict* | *TransferData* | *scnym_api(task='predict')* |
| **User** | *celltypist.annotate* | *LogisticRegression.predict* | *LinearSVC.predict* | *FindTransferAnchors*<br>*TransferData* | *scnym_api(task='train')*<br>*scnym_api(task='predict')* |

**B**

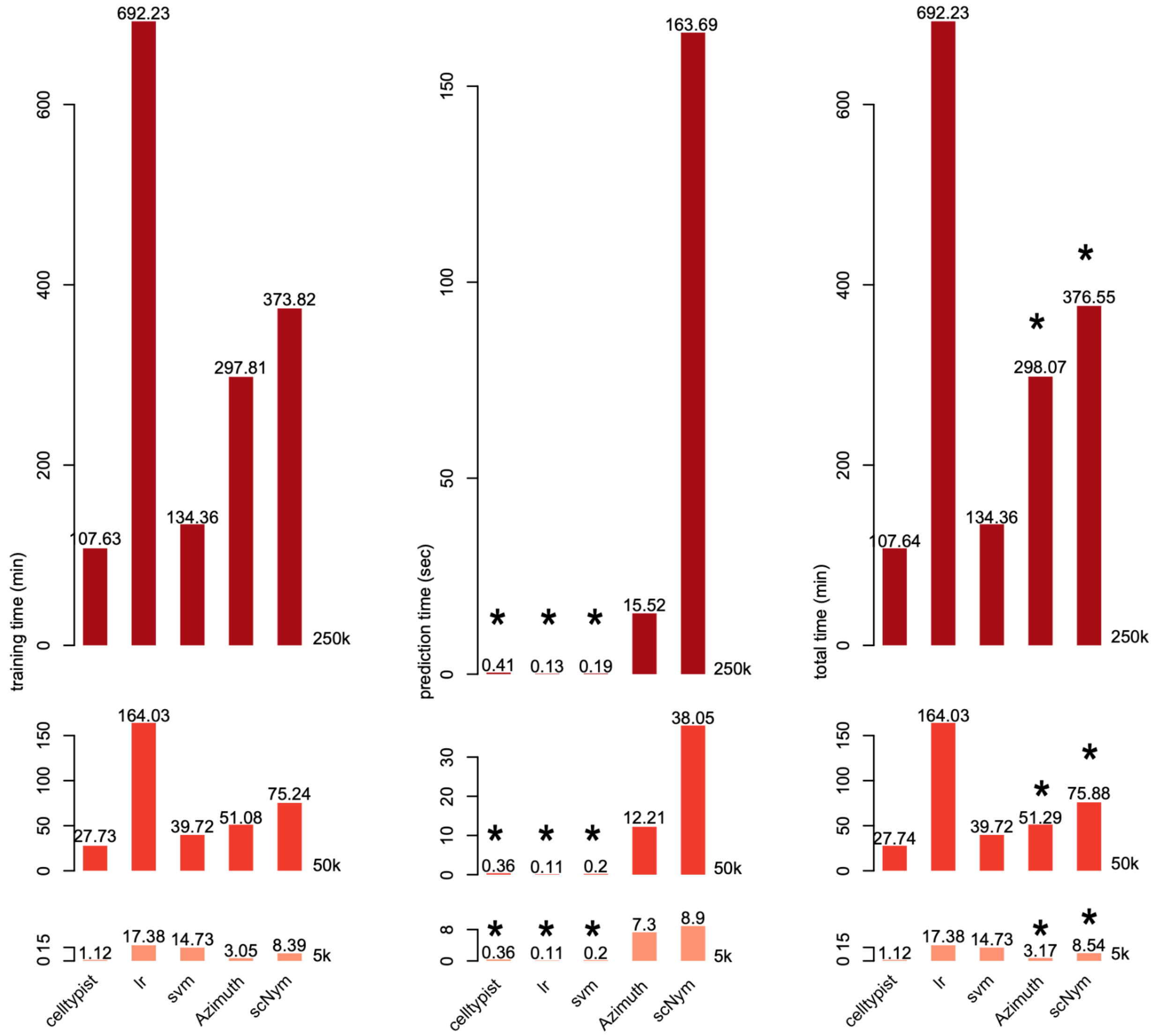

**Supplementary Note Figure 5**. **Benchmarking of CellTypist time complexity with other methods.** (**A**) Table summarising the split of different label transfer methods into the training and prediction steps. The "user" row shows the step/steps a user needs to get their prediction results after inputting the test data. (**B**) Bar plots showing the training time in minutes (left), prediction time in seconds (center) and total time in minutes (right) for the training dataset with 5,000 (lower), 50,000 (middle), and 250,000 (upper) cells, respectively. Five methods are assessed and the time is shown for each combination of training data and methods. Asterisks mark the user time for different methods.

**Supplementary Note Table 1**. Curated annotations for the low- and high-hierarchy models.

| High-hierarchy cell types | Low-hierarchy cell types |
|---|---|
| B cells | B cells |
| | Follicular B cells |
| | Germinal center B cells |
| | Memory B cells |
| | Naive B cells |
| | Transitional B cells |
| B-cell lineage | Immature B cells |
| | Pre-B cells |
| | Pre-pro-B cells |
| | Pro-B cells |
| Cycling cells | Cycling B cells |
| | Cycling DCs |
| | Cycling gamma-delta T cells |
| | Cycling monocytes |
| | Cycling NK cells |
| | Cycling T cells |

| | |
|---|---|
| DC | DC |
| | DC1 |
| | DC2 |
| | DC3 |
| | Migratory DCs |
| | Transitional DC |
| DC precursor | DC precursor |
| Double-negative thymocytes | Double-negative thymocytes |
| Double-negative thymocytes | Double-positive thymocytes |
| Early MK | Early MK |
| Endothelial cells | Endothelial cells |
| Epithelial cells | Epithelial cells |
| Erythrocytes | Erythrocytes |
| Erythroid | Early erythroid |
| | Late erythroid |
| | Mid erythroid |
| ETP | ETP |
| Fibroblasts | Fibroblasts |
| Granulocytes | Granulocytes |
| | Neutrophils |
| HCAImmune18_Blood_1 | HCAImmune18_Blood_1 |
| HCAImmune18_Blood_2 | HCAImmune18_Blood_2 |

| | |
|---|---|
| HCAImmune18_Blood_3 | HCAImmune18_Blood_3 |
| HCAImmune18_BoneMarrow_1 | HCAImmune18_BoneMarrow_1 |
| HCAImmune18_BoneMarrow_2 | HCAImmune18_BoneMarrow_2 |
| HSC/MPP | CMP |
| | Early lymphoid/T lymphoid |
| | ELP |
| | GMP |
| | HSC/MPP |
| | Megakaryocyte-erythroid-mast cell progenitor |
| | MEMP |
| | Neutrophil-myeloid progenitor |
| ILC | ILC |
| | ILC1 |
| | ILC2 |
| | ILC3 |
| | NK cells |
| | Transitional NK |
| ILC precursor | ILC precursor |

| | |
|---|---|
| Jaitin19_Omentum_1 | Jaitin19_Omentum_1 |
| Macrophages | Hofbauer cells |
| | Kidney-resident macrophages |
| | Kupffer cells |
| | Macrophages |
| Mast cells | Mast cells |
| Megakaryocyte precursor | Megakaryocyte precursor |
| Megakaryocytes/platelets | Megakaryocytes/platelets |
| MNP | MNP |
| Mono-mac | Mono-mac |
| Monocyte precursor | Monocyte precursor |
| Monocytes | Monocytes |
| Myelocytes | Myelocytes |
| pDC | pDC |
| pDC precursor | pDC precursor |
| Plasmablasts | Plasmablasts |
| Promyelocytes | Promyelocytes |
| T cells | CD8a/a |
| | CD8a/b(entry) |
| | Cytotoxic T cells |
| | Follicular helper T cells |
| | gamma-delta T cells |

| | |
|---|---|
| | Helper T cells |
| | MAIT cells |
| | Memory CD4+ cytotoxic T cells |
| | NKT cells |
| | Regulatory T cells |
| | T cells |
| | T(agonist) |
| | Tcm/Naive cytotoxic T cells |
| | Tcm/Naive helper T cells |
| | Tem/Effector cytotoxic T cells |
| | Tem/Effector helper T cells |
| | Tem/Effector helper T cells PD1+ |
| | Treg(diff) |
| | Type 1 helper T cells |
| | Type 17 helper T cells |
